# Dynamic Gene Regulatory Network Inference with Interpretable, Biophysically-Motivated Neural ODEs

**DOI:** 10.1101/2025.09.19.676870

**Authors:** Maggie Beheler-Amass, Chrisopher A Jackson, David Gresham, Richard Bonneau

**Affiliations:** Center for Genomics & Systems Biology, New York University, New York, NY; New York Genome Center, New York, NY; Genentech

## Abstract

Gene Regulatory Networks (GRNs) are complex dynamical systems that modulate gene expression and drive transitions between phenotypic cell states. Determining these networks is crucial in understanding how gene dysregulation can lead to phenotypic variation and perturbation responses. We present a novel biophysically-motivated neural ordinary differential equation (ODE) model framework with a biologically interpretable deep learning architecture that leverages dynamic single-cell data: in-CAHOOTTS (gene regulatory network Inference with Context Aware Hybrid neural-ODEs on Transcriptional Time-series Systems). Our approach combines accurate prediction with mechanistic interpretability, decomposing gene expression dynamics into fundamental biophysical processes (mRNA transcription and degradation) while inferring regulatory network structure. We validate in-CAHOOTTS by learning the *Saccharomyces cerevisiae* response to rapamycin treatment, and learning the dynamic cell cycle progression. The trained model accurately predicts gene expression trajectories and identifies biologically relevant rapamycin response regulators. The framework achieves stable long-term predictions with time frames extending 30 times beyond training data, maintaining realistic oscillatory dynamics for over 40 hours in cell cycle modeling, demonstrating that neural ODEs can capture true biological attractors rather than merely fitting data. We argue that interpretable neural ODEs can successfully model complex biological dynamics while revealing mechanistic insights essential for understanding living systems. By bridging extensible machine learning frameworks and mechanistic biology, in-CAHOOTTS represents a significant advance toward deep learning systems that both predict biological outcomes and explain the mechanisms driving them.

## 1 Introduction

Transcriptional Gene Regulatory Networks (GRNs) are dynamical systems that connect transcription factors to their target genes and drive cellular responses to stimuli, developmental transitions, and disease states (1; 2; 3; 4; 5). Uncovering these networks is crucial in understanding how gene dysregulation can lead to phenotypic variation and diseases. The goal of transcriptional GRN inference (GRNi) is to determine a network which represents how transcriptional activators and repressors control RNA transcription, to create a systems-level understanding of cellular regulation (6; 7; 8; 9; 10). Existing methods for GRNi often rely on statistical correlation rather than biologically motivated interactions, or rely on deep learning architectures that act as black boxes; these frameworks are difficult to interpret for biological mechanism (11; 12; 13).

Single-cell RNA sequencing (scRNA-seq) has been used to measure dynamic transcriptomic responses, providing time-series data that captures cellular responses to perturbations and transitions between phenotypic states (14; 15; 16; 17; 18; 19; 20; 21; 22; 23). However, computational methods struggle to fully exploit the temporal richness of scRNA-seq datasets while maintaining biological interpretability (24).

Deep learning frameworks excel at modeling complex interactions, but they typically discard central biological concepts such as sparsity and interpretability in favor of model complexity (25; 11; 26). These black-box approaches also lack the interpretability to uncover the causal mechanisms driving the dynamic biological systems, the underlying GRN (24; 1). Knowledge-primed neural networks offer a potential solution, where models learn from known biological interactions while maintaining interpretable parameters at each layer (9; 27; 28; 7; 10). By embedding known biophysical information into our models, determined from experimental assays, we can train knowledge-informed models that are biased towards true biological mechanism. This approach has been shown to improve GRNi accuracy by enriching true positives and providing causal grounding for inferred networks (9; 28; 7; 10; 29; 30; 19). These knowledge-informed approaches must also capture the inherently dynamic nature of gene regulation, where regulatory relationships evolve over time in response to cellular conditions and perturbations.

Neural ODEs represent a natural solution to these challenges. Complex dynamical systems can be modeled using ordinary differential equations (ODEs), making them ideal candidates for inferring dynamic GRNs (31). Deriving explicit physical models for each molecular nonlinear interaction is likely intractable due to the complexity of transcriptional regulation; in contrast, neural ODEs enable us to learn the regulatory network models using machine learning while capturing the underlying mathematical structure of biological processes in differential equation models of expression over time (32; 33; 14). Neural ODEs are particularly useful for interpolating predictions between missing data, predicting outcomes of biological experiments *in silico* from problems where training data may be sparse or sampled with an irregular time-axis (34; 35; 36). Neural ODEs provide a foundation for physics-informed neural networks (PINNs) in biology, where models can incorporate known biophysical constraints and differential equation structures to respect fundamental biophysical processes, making them ideal candidates for studying dynamical biological systems (37). Recent work suggests neural ODEs may show promise in predicting unseen temporal gene expression, predicting perturbation responses, and can be encoded to uncover GRNs while maintaining biological interpretability using physics-informed deep learning (28; 38; 37; 17; 19; 39; 19). We aim to enhance these models by implicitly learning biophysical parameters that control mRNA dynamics (40), which has been shown to improve GRNi (10; 41; 42; 43; 44).

Here, we present in-CAHOOTTS (gene regulatory network **In**ference with **C**ontext **A**ware **H**ybrid neural-**O**DEs **o**n **T**ranscriptional **T**ime-series **S**ystems), a novel biophysically-motivated Neural ODE framework that models the underlying biophysical processes of mRNA transcription and degradation. Our approach uses interpretable deep learning, constrained by prior biological knowledge, to infer GRNs while decomposing RNA velocity into its positive (mRNA transcriptional synthesis) and negative (mRNA degradation) components. This enables both accurate prediction of future cellular states and discovery of the regulatory networks governing these dynamics.

We demonstrate the effectiveness of in-CAHOOTTS using high-resolution time-series data from *Saccharomyces cerevisiae* responding to rapamycin treatment and undergoing cell cycle progression (10; 45). The model successfully predicts gene expression dynamics at unseen time points, identifies key transcription factors driving cellular responses, and maintains stable long-term predictions extending 30 times beyond training time frames. Our interpretable architecture reveals biological insights that would be invisible to black-box approaches, such as the disconnect between latent transcription factor activity and the observable mRNA expression of those transcription factors, and the underlying GRN driving the system.

## 2 Goals of Predictive Modeling

Understanding how gene regulatory networks (GRNs) drive cellular responses requires predictive models that capture the dynamics and causal relationships within biological systems. Our goal is to build predictive models where specific perturbations of the GRN yield observable phenotypic responses, enabling us to perform *in silico* experiments. Ideally, we also want to decompose the forces driving changes in the system into their underlying biophysical components. By starting the biological system in an initial state and predicting the phenotypic response to a perturbation over time, we can understand these changes in the context of separable biological processes (Figure 1A, top panel).

**Figure 1:**
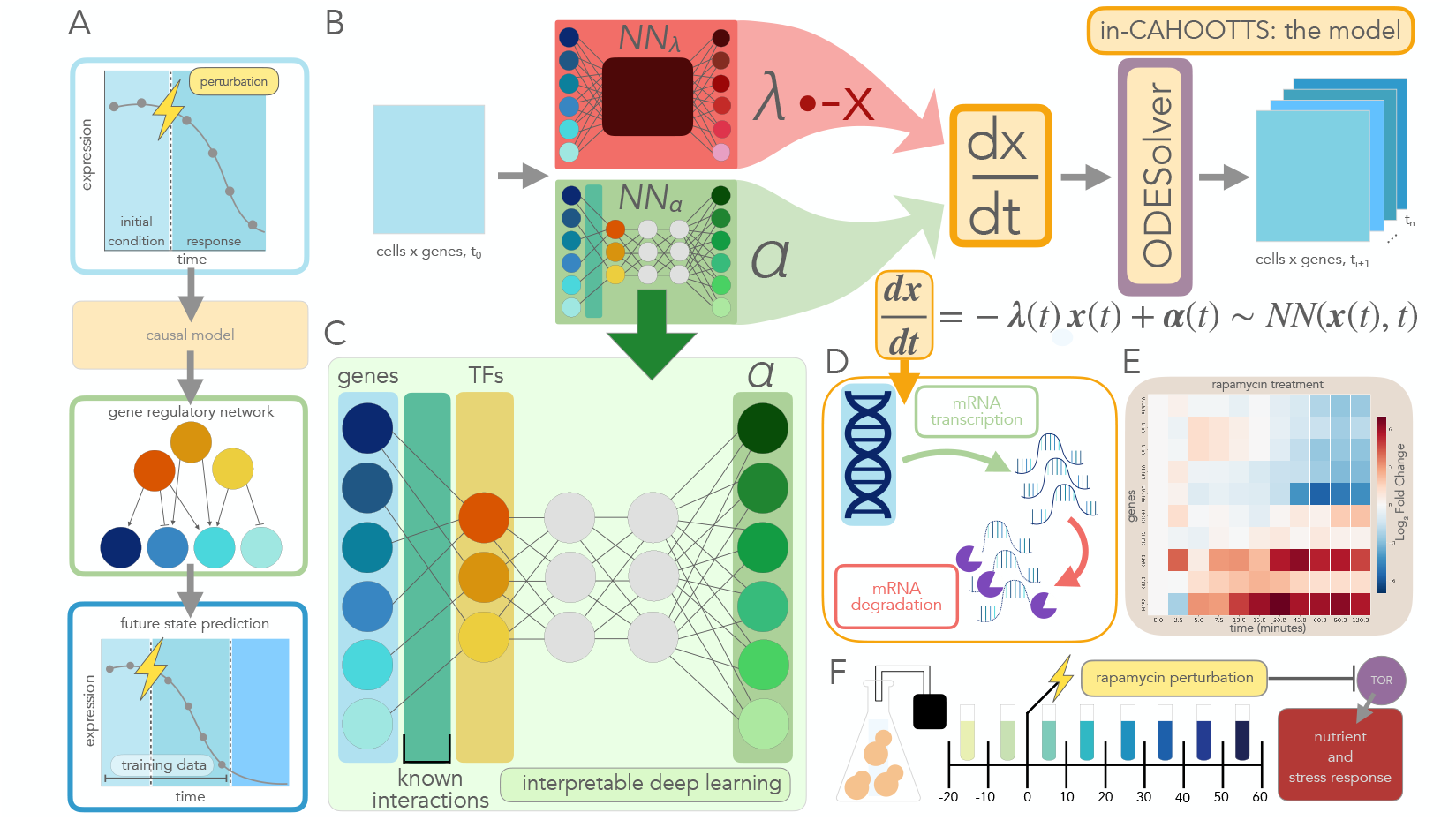
in-CAHOOTTS framework for dynamic, biophysically-motivated interpretable gene regulatory network inference. **(A)** Time-series gene expression data, collected from cells responding to a perturbation is used to train a causal model to infer the underlying GRN and predict future gene expression at unseen time points. **(B)** The in-CAHOOTTS neural ODE architecture takes gene expression *x* across cells at an initial condition *t*_0_ and predicts RNA Velocity (^*dx*^*/*_*dt*_) from a positive-component (*NN*_*α*_) and a negative-component (*NN*_*λ*_) neural network. *NN*_*α*_ estimates mRNA transcription rates on a per gene basis, and can be interpreted internally as a gene regulatory network. *NN*_*λ*_ is a black-box feed-forward network whose output *λ* is an estimate of mRNA decay rate on a per gene basis. Outputs of the both *NN*_*α*_ and *NN*_*λ*_ are constrained to be positive. *λ* is multiplied by −*x* to estimate mRNA degradation and is combined with *α* to estimate ^*dx*^*/*_*dt*_, (Equation 14) which is fed into an ODESolver to predict gene expression at future time points, and the model is trained by comparing predictions to observed future time points. **(C)** Transcription rate prediction model *NN*_*α*_ is interpretable as a gene regulatory network. The input gene layer is connected to a latent space that is interpretable as transcription factors by constraining encoder weights with an adjacency matrix of prior known transcription factor-gene interactions. The underlying GRN is inferred by analyzing the importance of TF-gene connections learned by the model. **(D)** Gene expression change 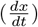 is the combination of positive mRNA transcriptional pathways, and negative mRNA decay pathways, both of which are tightly regulated by cell signaling pathways. **(E)** *S. cerevisiae* gene expression changes from bulk RNA sequencing after rapamycin treatment for 10 selected genes. Downregulated genes on top are ribosomal genes, and upregulated genes on the bottom are stress response. **(F)** Exponentially dividing budding yeast (*S. cerevisiae*) cultures are grown for 20 minutes. The culture is then perturbed by adding the TOR inhibitor rapamycin, and sampled for an additional 60 minutes. Cultures are sampled continuously by peristaltic pump into an excess of saturated ammonium sulfate, and sequenced by single-cell RNA sequencing to get a continuous trajectory of cells during perturbation response (10).

Our first objective is to build a model that takes dynamic gene expression data and learns how cells respond to perturbations, capturing both the magnitude and temporal dynamics of gene expression changes. Our second objective is to interpret this model in a biological context, to uncover the underlying dynamical system driving these responses; for the transcriptome, this is the transcriptional GRN consisting of transcription factors regulating target genes (Figure 1A, middle panel). However, rather than treating gene expression changes as black boxes, we want to separate the underlying biophysical processes—separating transcription of mRNA from mRNA degradation, which together drive observed dynamics (Figure 1D). Once we establish this model framework, we can use it to predict future states of the system: for example, how will cells continue responding to perturbations at time points beyond our training data (Figure 1A, bottom panel)? Finally, we want models capable of long-term predictions, like the complex oscillatory behavior of the cell cycle. This predictive capability can potentially extend to counterfactual scenarios; if we start from a different initial state, such as a diseased cell treated with the same perturbation used on healthy cells, can we forecast the temporal response?

With these predictive biology goals in mind, we developed **in-CAHOOTTS**: gene regulatory network **In**ference with **C**ontext **A**ware **H**ybrid neural-**O**DEs on **T**ranscriptional **T**ime-series **S**ystems.

## 3 The in-CAHOOTTS Model

The time derivative (^*dx*^*/*_*dt*_) of gene expression (*x*), can be interpreted as instantaneous RNA velocity, the rate of change in abundance over time of an mRNA transcript. RNA velocity has been used to infer directed differentiation trajectories from transcriptomic data and infer GRNs (14; 15; 17; 46; 47). The in-CAHOOTTS model is designed to implicitly estimate RNA velocity, the local derivative of expression (^*dx*^*/*_*dt*_), from gene expression *x*. The estimated RNA velocity from 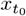, gene expression at the initial time point, is fed into a Neural ODESolver to predict gene expression across future time points 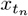 for various experimentally-measured trajectories (Figure 1B).

Following the formulation proposed in (10), our model represents gene expression dynamics through a first-order ordinary differential equation (Equation 1):

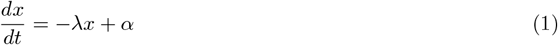

The positive, transcriptional component *α* is estimated by an interpretable neural network based on an architecture (9) where connections in the transcriptional model can be interpreted as a GRN (Figure 1C, top panel). The architecture incorporates a context-specific matrix of prior known interactions between transcription factor proteins and their target genes to regularize and constrain the neural ODE solutions to biologically real interactions. The prior acts as a biologically-informed soft constraint on the Neural ODE, guiding the model towards established regulatory mechanisms while still allowing for data-driven discovery of novel interactions. The mRNA decay rate *λ* is estimated by a neural network, and *λ* is multiplied by gene expression *x*_*t*_ to obtain an estimate of the negative, degradation component of velocity. Both *α* and *λ* can vary over time and are modeled separately for each gene. RNA Velocity (^*dx*^*/*_*dt*_) is modeled as the combination of transcription and degradation (48; 49), which allows the model to separate these two different types of mRNA regulation (Figure 1D).

The separate neural networks are trained together, and the combined model estimates gene expression at future time points. By training on expression trajectories, and by constraining the transcriptional component of the model for interpretability, this framework both predicts future gene expression and uncovers the gene regulatory network driving the overall dynamics of the system.

### 3.1 Neural Ordinary Differential Equations

Neural ODEs are formulated as (32; 33) (Equation 2):

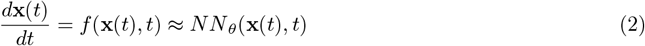

where the time derivative, or velocity, is governed by a set of ODEs constrained by the underlying GRN, and *f* is a vector field function of the instantaneous cell states, estimated using a neural network with learnable parameters *θ* to yield *NN*_*θ*_ (32; 28). Starting from an initial condition *x*(*t*_0_), we can estimate an output *x*(*t*_*n*_) to be the solution to the ODE initial value problem at time *t*_*n*_ via a numerical ODESolver with an adaptive step size:

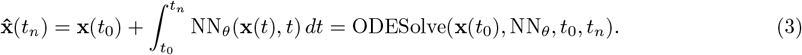

The loss function is then:

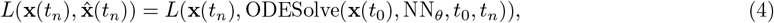

where *x*(*t*_0_) is used to predict time point *x*(*t*_*n*_), and mean squared-error is used to compare the actual versus predicted output. Gradients are calculated via the loss functions using the adjoint sensitivity method, computing derivatives for back-propagation through the Runge–Kutta (rk4) ODESolver (32).

### 3.2 Model Framework and Architecture

To adapt the Neural ODE (2) for this framework for dynamic GRNi we model mRNA decay constant (*λ*) and transcriptional output (*α*) (Equation 1) as time-dependent parameters which are estimated by learned functions *f*_*λ*_ and *f*_*α*_:

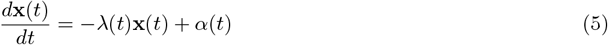

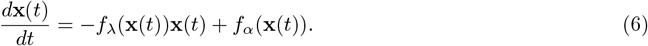

RNA velocity (^*dx*^*/*_*dt*_), the mRNA decay constant (*λ*) and the transcriptional output (*α*) are implicitly learned by interpreting the neural ODE outputs for *f*_*λ*_ and *f*_*α*_ in Equation 6 as estimates of *λ* and *α* in Equation 5 on a per gene basis (Figure 1B). Model predictions are constrained to be strictly positive by nonlinear activation functions, GeLU (50), RELU (51) and Softplus (52), to maintain biological interpretability while learning the contributions of degradation and transcription driving the changes in gene expression.

#### 3.2.1 Estimating mRNA Transcription

The architecture for *f*_*α*_ is based on an interpretable deep learning architecture for transcriptional GRN inference (9) (Supplemental Figure S1; Equations 7-10). The input expression data **X** (*N*_genes_ = 5726) at *t*_0_ is encoded into a latent space **h**^(TF)^ with dimensionality equal to the number of transcription factors (*N*_tfs_ = 158), allowing the first hidden layer of the latent space to be interpretable as transcription factor (TF) activity (see Prior Network Knowledge). The ranked connections (see Section 6.5.2) between **h**^(TF)^ nodes in the latent space, i.e. the TFs, and the target genes in the outer layer are therefore interpretable as an inferred GRN. The TF layer is constrained to be positive by the ReLU activation function, making the embeddings in the latent layer biologically interpretable as transcription factor activity (Figure 1C).

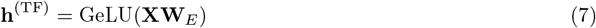

The TF layer is linearly embedded into two intermediate layers with fully connected weights and the GeLU activation function is applied after each embedding.

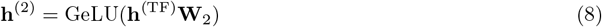

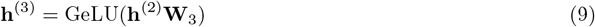

**h**^(3)^ is decoded from the hidden layers to the output transcriptional rates per gene using fully connected weights **W**_*D*_, and the ReLU activation is applied to constrain the outputs to be strictly positive, allowing the output layer to be biologically interpretable as a rate of transcriptional output per gene *α* (Figure 1C).

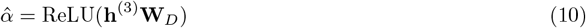

#### 3.2.2 Incorporating Prior Network Knowledge

The prior in this framework constrains our model to known transcriptional regulatory interactions, helping the neural ODE find the most biologically relevant solution. Applying known regulatory relationships as a prior has been previously shown to allow GRN inference with neural ODEs (28).

Here, prior knowledge is represented as a sparse adjacency matrix *P*, where edges *p* are boolean (*p* ∈ {0, 1}), that connects 158 TFs to 5726 genes (Figure 1C). The prior is derived from documented transcription factor-gene interactions (53). A hand-curated gold standard regulatory network (29), which is a subset of the prior *P*, contains 1403 highest-confidence edges connecting 98 TFs to 993 genes.

The prior *P* is applied to the transcriptional model by constraining *f*_*α*_ model weights **W**_*E*_ (Equation 7); *P* and **W**_*E*_ share the same dimensionality (*N*_genes_ x *N*_tfs_). Initial weights for **W**_*E*_ are set such that edges that are zero in *P* are also zero in **W**_*E*_.

During training, we allow all edges in **W**_*E*_ to vary from zero, but apply a regularization term that penalizes deviation from the prior knowledge:

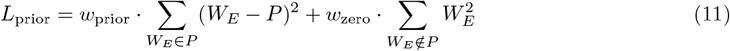

where *W*_*E*_ ∈ *P* are edges in the prior *P, W*_*E*_ ∉ *P* are edges not in the prior *P*, and weights *w*_prior_ and *w*_zero_ are fixed regularization coefficients. This approach encourages the model to preserve documented interactions while allowing data-driven discovery of novel connections (Figure 1C).

#### 3.2.3 Estimating mRNA Decay

The architecture for *f*_*λ*_ is based on a black-box feed-forward network (Supplemental Figure S2; Equations 12-13). Expression (*N*_genes_) for each cell (observation) at *t*_0_ is linearly embedded into a latent space (with dimensionality *N*_tfs_) using fully connected weights and the Softplus activation function is applied. The hidden layer is decoded using fully connected weights and the Softplus activation is applied to yield an estimated decay rate per gene.

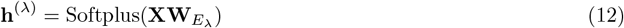

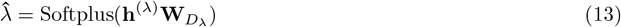

#### 3.2.4 Inferring RNA Velocity and Predicting Expression

The outputs from the transcription and decay models are combined to estimate RNA Velocity at time *t*. Decay rate estimate 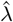 is multiplied by -1 to represent degradation of mRNA, and multiplied by RNA expression to estimate the negative component of RNA velocity.

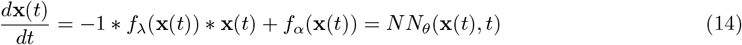

Expression data **x**(*t*_0_) are fed into the Transcription and Decay architectures via the ODESolver, which calls the forward function within the base model architecture. The ODESolver gives the model an initial value at a starting timepoint, and the model outputs an inferred value for RNA Velocity, 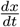. The velocity is then inferred across each time step given to the ODESolver, and *X*_*t*_ is predicted for each time point given to the ODESolver (Figure 1B).

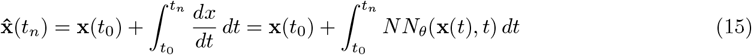

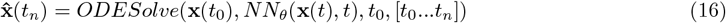

The initial condition is the only information embedded into the model on the forward pass. The neural ODE samples the vector field it learns at each of these time points in the solution space and uses an iterative Runge–Kutta (rk4) ODESolver to predict the next time points (Figure 1B). Loss (Equation 4) is calculated as mean-squared error between **x**(*t*_*n*_) and 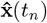, embedding information from later time points during model weight updates in the backward pass (32).

## 4 Results

### 4.1 Dataset characteristics and experimental design

We train the in-CAHOOTTS model on a pre-existing single-cell *Saccharomyces cerevisiae* dataset from Jackson et al. (10), where cells are responding to a rapamycin perturbation. Rapamycin is an inhibitor of the nutrient-sensing Target of Rapamycin (TOR) pathway, causing exponentially-growing yeast cells in rich media to shift from a proliferative transcriptional state to a nitrogen starvation state, upregulating metabolic and stress response transcripts, and downregulating ribosomal transcripts (Figure 1E, Supplemental Figure S3). In the original experimental design, the culture was continually pumped into an excess of saturated ammonium sulfate, which fixes the transcriptome and maximizes the time resolution of the transcriptional response. After collecting 20 minutes of exponentially-dividing cells, the culture was perturbed by adding rapamycin, and then an additional 60 minutes of cells were collected post-perturbation. The total time-series dataset contains 173,361 cells continuously sampled over 80 minutes across 8 time pools, in biological duplicate (Figure 1F) (10).

For training, cells are arranged into trajectories using pre-computed temporal assignments from Jackson et al. (10), with each trajectory containing 60 cells sampled one minute apart (Figure 2A). Cells at t=0 from the trajectories serve as initial conditions for the neural ODE, which then predicts gene expression for the subsequent 60 time points (minutes 1-60) (Figure 2A). The neural ODE uses the first cell at t=0 (from the pool of untreated cells) as the initial condition, and then predicts gene expression for the next 60 time points (minutes 1-60) (Figure 2A). Predicted gene expression 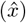 in the trajectory is compared to observed gene expression in the trajectory (*x*), and the mean squared error is backpropagated through the model for training. Trajectories were randomly resampled after every training epoch.

**Figure 2:**
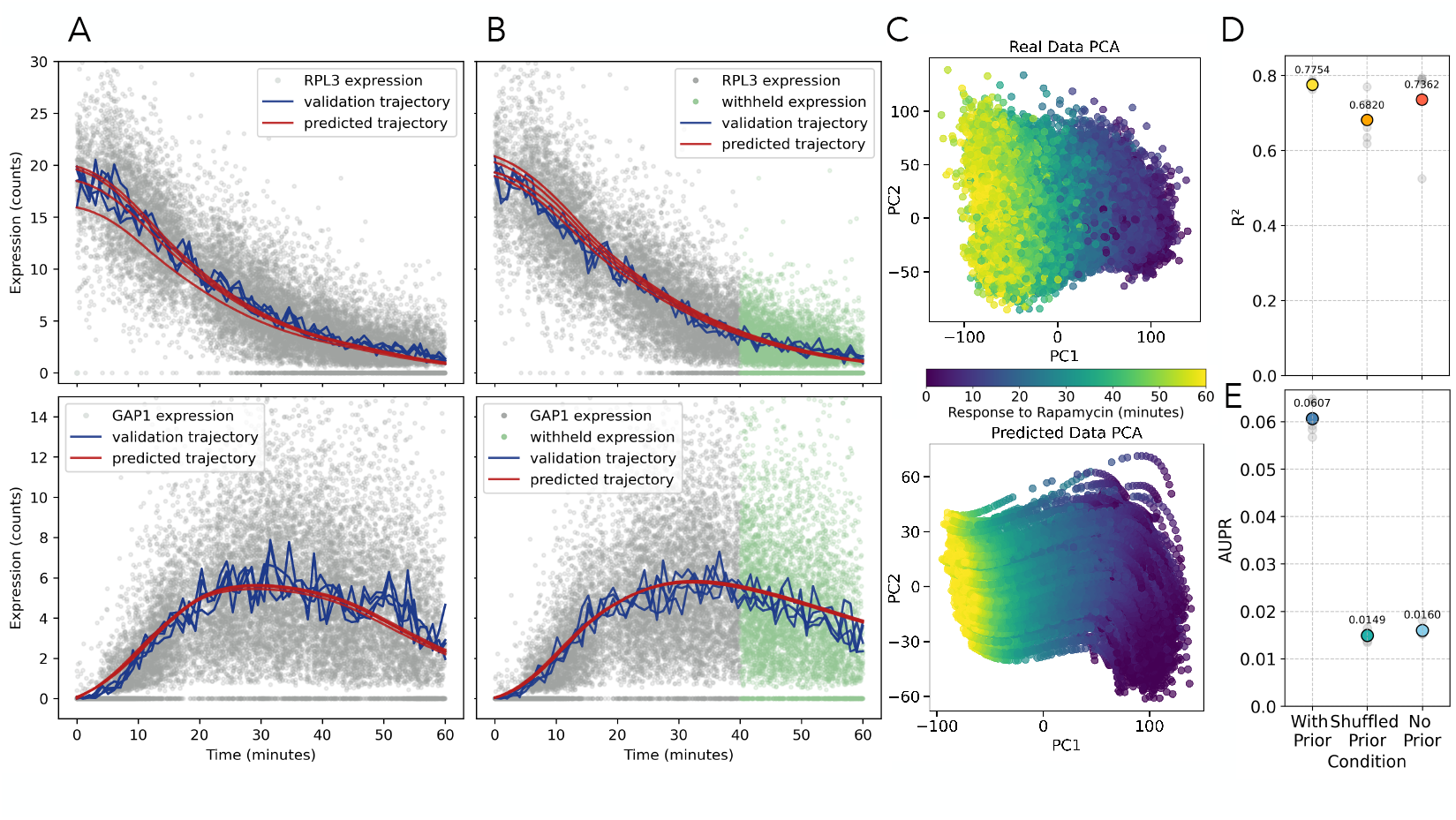
in-CAHOOTTS predicts *S. cerevisiae* gene expression dynamics in response to rapamycin perturbation and recovers the underlying gene regulatory network. **(A)** Predicted expression from in-CAHOOTTS model trained on data from t=0 to t=60 of ribosomal protein gene *RPL3* (top) and amino-acid transporter gene *GAP1* (bottom). Gray dots are individual validation cells (*n* = 19800); blue lines are the average of 20 cells from 1 minute windows, assembled into a validation trajectory (t=0 to t=60 minutes). Red lines are predicted trajectories (t=0 to t=60 minutes at 1 minute intervals). **(B)** Predicted expression from in-CAHOOTTS model trained on data from t=0 to t=40 of *RPL3* and *GAP1* expression, plotted as in **A**, with cells from t=40 to t=60 (not included in training data) colored in green **(C)** Principal component plot of the first two PCs with actual cells colored by assigned times (top), compared to predicted expression colored by time on the prediction trajectory (bottom) **(D)** Model predictive performance, quantified as *R*^2^, on models trained with a biological prior matrix, a shuffled (random) biological prior matrix, and no biological prior. Gray dots are individual values for a 5-fold crossvalidation, and colored dots are the mean. **(E)** Model GRN reconstruction performance, quantified as AUPR, on models trained with a biological prior matrix, a shuffled (random) biological prior matrix, and no biological prior. Scoring is done on genes held out of the prior matrix by cross-validation, gray dots are individual values for each cross-validation, and colored dots are the mean.

The in-CAHOOTTS model predicts downregulated expression of ribosomal genes, as with ribosomal protein *RPL3*, and predicts upregulation of nutrient stress response genes, as with amino acid transporter *GAP1* (Figure 2A), (see *R*^2^ value in Figure 2D) (54). To show that this is not purely memorization, we trained a in-CAHOOTTS model using trajectories from 0-40 minutes, with no information about expression from minutes 41-60, and then used that trained model to predict expression from t=0 to t=60. This model is also able to predict gene expression trajectories very well, and accurately predicts expression dynamics from t=40 to t=60. As a specific example, GAP1 expression is predicted to decrease between timepoints 40-60, matching the observed data which was not used during training.

To further validate the in-CAHOOTTS model’s ability to make out-of-training future predictions, we predict expression to t=120 minutes, which is double the length of the training time, and compare these predictions against comparable rapamycin response bulk RNA-seq expression data (Supplemental Figure S5) (10). These future predictions are accurate, successfully capturing both the magnitude and dynamics of key rapamycin-responsive genes; there correctly predicted continued downregulation of ribosomal genes (*RPL3*), sustained upregulation of stress response factors (*GCN4*), and the activation and eventual repression of nutrient response genes *GAP1* and *MEP2*. This validation demonstrates the in-CAHOOTTS model’s ability to learn generalizable biological dynamics, enabling robust extrapolation to unseen experimental conditions and timeframes.

Finally, we explored the limits of the framework when data is more sparsely sampled (Supplemental Figure S7). We subsampled the 60-minute trajectories at intervals of 30, 20, 10, and 5 minutes and compared performance against the original 1-minute sampling method. *R*^2^ values remain relatively high across these sampling densities when using the prior, suggesting the model can make accurate predictions on datasets less densely sampled than ours and can smoothly interpolate to predict expression at unseen time points. However, the model fails to converge when trained with 30-minute intervals, indicating limitations in the current formulation for truly sparse temporal datasets.

While we show *RPL3* and *GAP1* as specific examples of down- and up-regulation, the trained model has high performance to predict the entire transcriptome. This can be seen by comparing the principal component (PC) plot of the first two PCs for the single-cell transcriptomes (Figure 2C, top) with the first two PCs for the predicted trajectories (Figure 2C, bottom). The in-CAHOOTTS predicted transcriptomes show a smooth trajectory advancing through time, which is qualitatively comparable to the overall dynamics of the rapamycin response in the observed data. We do note that the model predictions show reduced variance compared to actual data, visible as decreased range in PC2, reflecting the deterministic nature of the learned dynamics, which does not incorporate either biological or sampling noise. Quantitatively, the trained model has an *R*^2^ during cross-validation of 0.775 (Figure 2D), comparing predicted trajectories from t=1 to t=60 to validation trajectories, indicating that the model is generally successful at making transcriptome-wide predictions.

In addition to predicting transcriptomes, the in-CAHOOTTS model also attempts to reconstruct a GRN which reflects biological TF to gene regulatory relationships. We initially train the model for 500 epochs without a prior, allowing the nODE to learn cellular dynamics from a data-driven approach. However, without prior knowledge, the model does not learn biological connections between transcription factor proteins and their target genes (Figure 2D). We introduce prior knowledge about TF to gene regulation by constraining the transcriptional encoder model, penalizing encoder weights which are not in the prior matrix (*w*_*prior*_ = 0.99 *w*_*zero*_ = 0.5), and train the model for an additional 600 epochs. We quantify success at reconstructing biological regulation in the in-CAHOOTTS model by calculating the area under the precision-recall curve for genes which are withheld entirely from the prior knowledge matrix (Figure 2E), comparing the fully trained models with the prior to the trained models without a prior, and to models where the prior was randomly generated by shuffling. Gene expression predictions (quantified as *R*^2^) perform similarly with or without prior knowledge, but interpretability for GRN reconstruction requires known prior relationships between TFs and genes.

We compare the in-CAHOOTTS model against an out of the box, non-fine-tuned Neural ODE baseline with comparable depth and parameters. This baseline compares to Neural ODE that lacks biophysical decomposition, interpretable architecture, and prior biological knowledge. This comparison highlights the critical importance of our methodological innovations: in-CAHOOTTS achieves 0.775 R2 compared to 0.483 for vanilla Neural ODE (60 percent improvement in predictive accuracy) and 0.0607 AUPR versus 0.0174 (3.5-fold improvement in regulatory network recovery) (Supplemental Figure S9). These substantial im-provements across both prediction and network inference demonstrate that the added complexity of our biophysical framework is not only justified but necessary for capturing meaningful biological dynamics.

### 4.2 Decomposing gene expression predictions into biophysical estimates of RNA degradation and transcription

The derivative of gene expression (^*dx*^*/*_*dt*_ or RNA velocity) is implicitly estimated by the Neural ODE and used by the ODESolver to predict future expression. We compare the in-CAHOOTTS predictions for ^*dx*^*/*_*dt*_ against empirical estimates for ^*dx*^*/*_*dt*_ for each cell generated by regressing gene expression against time within a local neighborhood of cells (10). These empirical estimates, owing to the continuous sampling method used, are viable as a baseline comparison for model output.

The in-CAHOOTTS velocity predictions match local empirical estimates for both *RPL3* and *GAP1*, though the model slightly overestimates the initial velocity at t=0 for *GAP1*, where expression begins at zero (Figure 3A). in-CAHOOTTS predictions of RNA Velocity are estimated by combining the output of *NN*_*α*_ and −*x* * *NN*_*λ*_ (Equations 5 and 6). We can interpret these velocity components before combination (Figure 3B), in order to study separate decay and transcription mechanisms.

**Figure 3:**
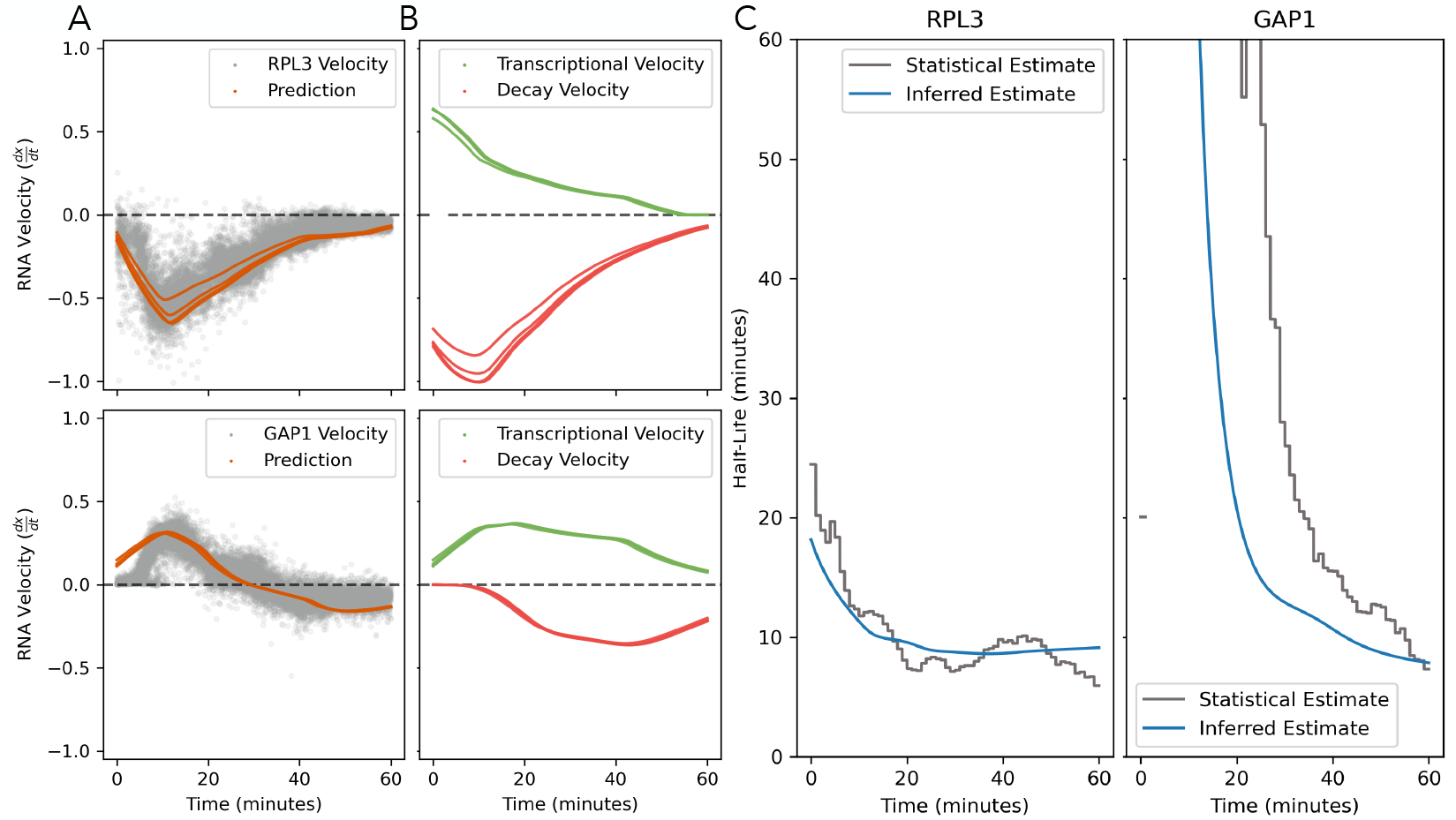
in-CAHOOTTS decomposes RNA velocity into biophysical estimates of decay and transcription. **(A)** Gray dots are estimates of RNA velocity (^*dx*^*/*_*dt*_) for each cell held out as validation, generated by regressing gene expression *x* against time *t* within a local graph neighborhood. Orange lines are in-CAHOOTTS predicted velocities plotted as trajectories. Ribosomal protein gene *RPL3* (top) and amino acid permease gene *GAP1* (bottom) are separately plotted. **(B)** in-CAHOOTTS predicted velocities, separated into green mRNA transcription trajectories (*N N*_*α*_ or 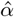 and red mRNA degradation trajectories (− \**x NN*_*λ*_ or 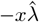). **(C)** Gray lines show time-dependent constrained optimization estimate of RNA decay rate from experimental data, plotted as half-life (*t*_1*/*2_ = ^*log*(2)^*/*_*λ*_). Blue lines are in-CAHOOTTS predicted decay rate trajectories 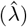 plotted as half-life.

### 4.3 mRNA degradation rate estimation

The negative biophysical component of RNA velocity, RNA degradation, is a regulated biological process that can respond to perturbation resulting in substantive changes in RNA decay rate. in-CAHOOTTS estimates RNA decay rates 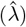 over time as the output of *NN*_*λ*_, and calculates RNA degradation as −*x* * *NN*_*λ*_ (Figure 3B). We compare the in-CAHOOTTS decay rate estimates to those generated from a sliding-bwindow constrained optimization method (10), which is viable owing to the continuous sampling method used to generate the data. We plot these values for comparison as RNA half-life in minutes (Figure 3C), and find similar patterns. The inferred RNA half-life for *RPL3* matches quite well with the statistical estimate. Initially, at t=0, *RPL3* transcripts have a half-life of 20 minutes; after perturbation, this transcript destabilizes and decays with a half-life of 10 minutes. *GAP1* expression is extremely low at t=0, and so the model predicts a low RNA decay rate (leading to a very large half-life in minutes). As *GAP1* expression increases, the half-life of the transcript decreases. These half-lives are consistent with findings from metabolic labeling studies, although experimental limitations preclude collecting RNA decay measurements with a high time resolution across the cell cycle (or any analogous cell state trajectory). (55).

### 4.4 Gene Regulatory Network Reconstruction

The positive biophysical component of RNA velocity, RNA transcription, involves the production of new RNA molecules that are regulated by TFs. and the production of new RNA molecules are regulated by TFs. in-CAHOOTTS estimates RNA transcription rates 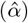 over time as the output of *NN*_*α*_ (Figure 3B). The interpretable architecture of the transcriptional model *NN*_*α*_ constrains encoder weights between genes and transcription factors using an adjacency matrix of prior known interactions to uncover interpretable connections to TFs driving variance in transcription. The architecture of *NN*_*α*_ uses an interpretable deep learning framework developed in SupirFactor, where the input layer is interpreted as genes, the latent space is interpreted as transcription factors and their activities, and the decoder, i.e. the connections between the latent space and the outer layer are interpretable as the underlying gene regulatory network (9).

The first latent layer **h**^(*T F*)^ of this model are embeddings that can be interpreted as transcription factor activities (TFA). TFA is a representation of the latent biological state of TFs, which quantifies the ability of that TF to change RNA transcription. This is distinct from the expression of TFs, as many TFs are regulated post-transcriptionally or post-translationally by signaling pathways, and predicted latent TFA determines which transcription factors are responding to rapamycin to cause expression changes.

We can further determine the regulatory network connections between TFs and target genes by explained relative variance analysis. This approach calculates the ratio of *R*^2^ for the full model to *R*^2^ for a partial model, where a TF node has been removed, to quantify how that TF is influencing gene expression for each gene. The percent of explained variance for each TF is interpreted as regulatory interaction strength in the gene regulatory network. Our architecture allows us to connect biophysical parameter estimation to reconstructing the regulatory mechanisms driving cellular responses.

Performing an elbow analysis on ranked ERV per connection in the inferred GRN, we found that most variance in this rapamycin response data is captured within the top 116 connections, encompassing 15 unique transcription factors (Supplemental Figure S10). This analysis was constrained to interactions present in the gold standard network to ensure biological validity. The top 5 regulators from this subset are the general amino acid control TF *GCN4*, the nutrient-limitation transcription factor *STE12*, the multifunction TFs *CBF1* and *ABF1*, and the stress response TF *MSN2* (Figure 4A), which align with known rapamycin response pathways.

**Figure 4:**
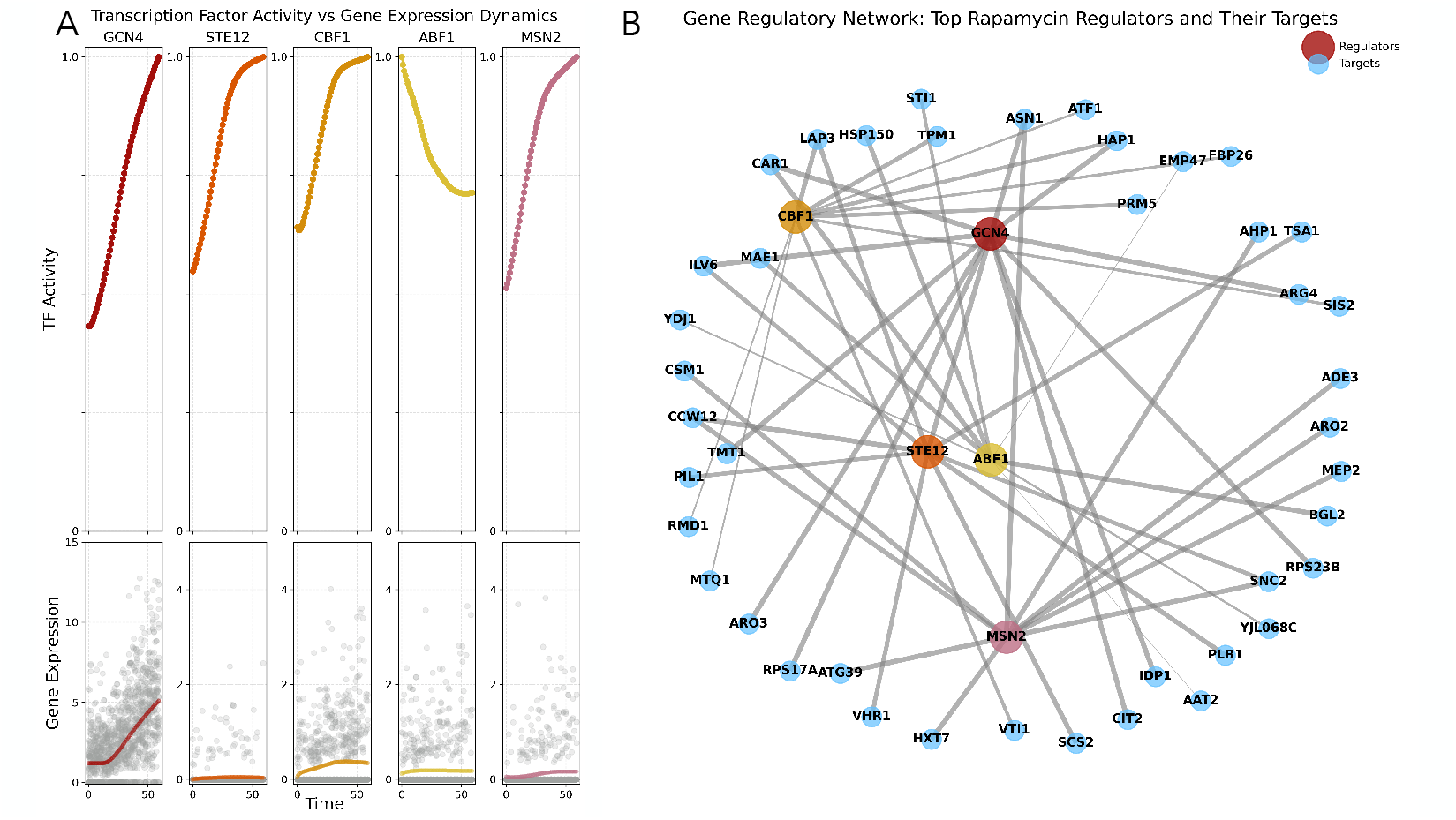
in-CAHOOTTS infers Transcription Factor Activities and reconstructs a Gene Regulatory Network. **(A)** Transcription factor activity predictions: Time-course dynamics of regulatory TFs identified by the model. The top panels show TF activities over time, revealing distinct temporal patterns (GCN4 and MSN2 increasing, STE12 and CBF1 decreasing, ABF1 stable). The bottom panels show corresponding gene expression levels, demonstrating that TF activity does not automatically correlate with gene expression. **(B)** Inferred gene regulatory network: Network structure showing the top rapamycin-responsive regulators (warm colors) and their target genes (cool colors). Edge connections represent high-confidence regulatory relationships learned by the model, demonstrating the regulatory architecture underlying the stress response. The network reveals hub transcription factors (GCN4, CBF1, ABF1, STE12, MSN2) that coordinate the cellular response to rapamycin treatment.

*GCN4* is the master regulator of amino acid starvation response and shows dramatic activity increases throughout the time course, consistent with TOR inhibition triggering a nitrogen starvation response. *GCN4* activity increases immediately on rapamycin treatment, *GCN4* mRNA expression is stable for 20 minutes, at which time it begins a modest upregulation, which is consistent with the known transcriptional and post-transcriptional regulation of the *GCN4* transcript (56). *MSN2, CBF1* and *STE12* activity also increase immediately on rapamycin treatment, although without any upregulation of their respective mRNA levels, indicating that their change in activity is linked to changes in post-transcriptonal regulation. Conversely, *ABF1*, which regulates ribosomal protein synthesis, shows declining activity, aligning with the expected downregulation of ribosomal gene transcription in response to rapamycin treatment (57).

The inferred GRN reveals the regulatory architecture of the dynamical system underlying this coordinated response (Figure 4B). The model identifies candidate regulatory connections between key stress-responsive TFs and their predicted target genes. For example, the framework predicts connections between GCN4 and amino acid biosynthesis genes, consistent with known amino acid starvation responses triggered by TOR inhibition. Similarly, predicted MSN2 connections to autophagy-related genes align with expected metabolic shifts from anabolic to catabolic processes. These predicted network relationships provide testable hypotheses about how transcriptional regulation coordinates cellular responses to rapamycin treatment.

### 4.5 Longer Time-Scale Predictions

We expect the in-CAHOOTTS model architecture to be particularly suited to making long-term future predictions. However, the long-term effects of rapamycin treatment are likely to be monotonic processes: the accumulation of mutations which facilitate rapamycin response escape, a general shift into a quiescent state, or sporulation into dormant ascospores. These mechanisms are not present in the training data and represent unidirectional cellular transitions that would eventually move the system beyond the learned dynamics. However, the training data contains a dynamic response to rapamycin and a dynamic response to progression through the cell cycle. In contrast, the cell cycle represents nonmonotonic, oscillatory behavior where cells repeatedly traverse the same sequence of transcriptional states (Figure 5A). We reason that the in-CAHOOTTS model is well-suited to making long-term predictions of gene expression during cell cycling because this cyclical process maintains cells within a defined attractor in transcriptional space. This allowed us to evaluate long-term predictions of oscillatory behavior in dynamic equilibrium.

**Figure 5:**
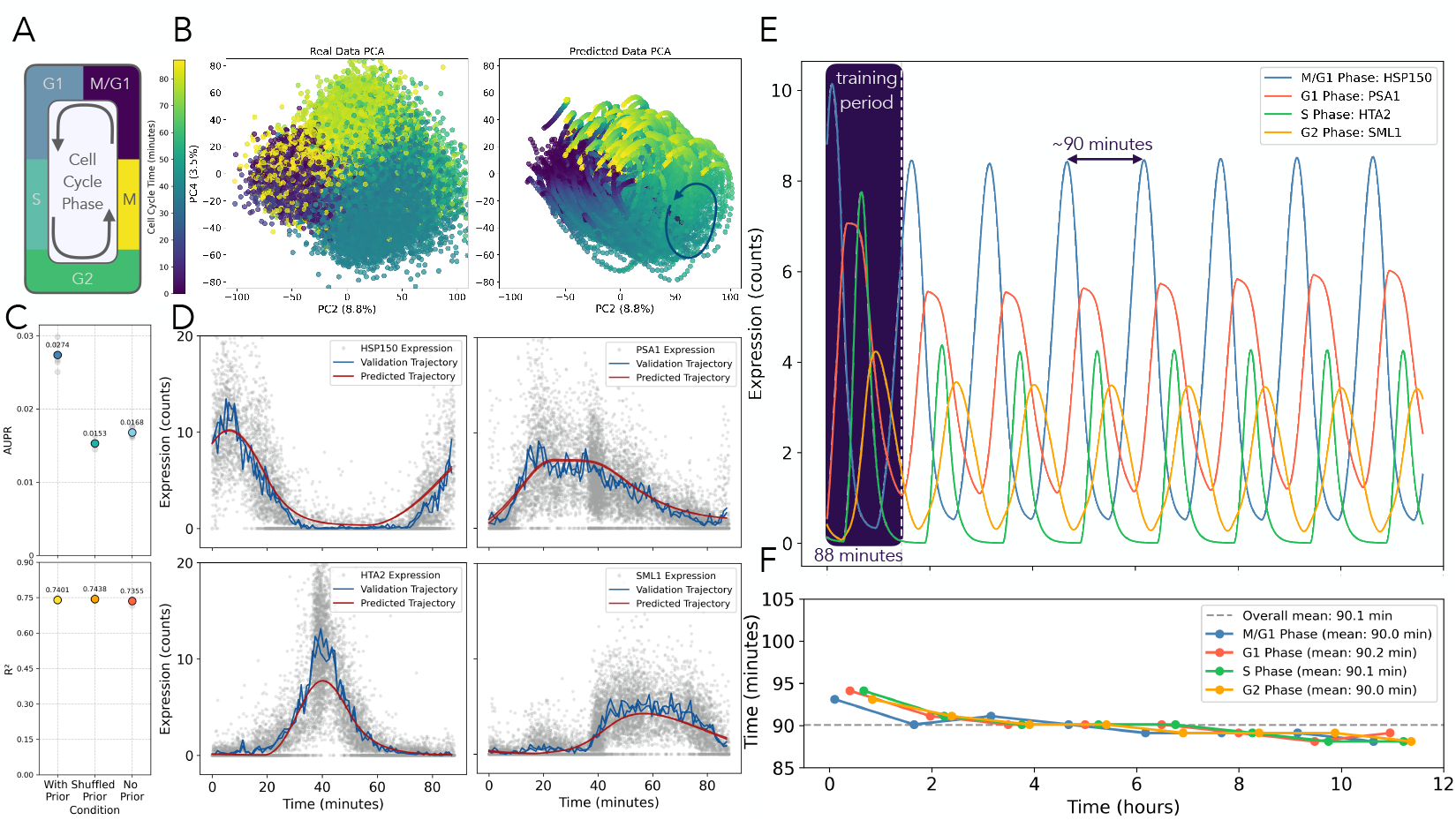
in-CAHOOTTS predicts S. cerevisiae cell cycle dynamics and enables long-term oscillatory predictions. **(A)** Schematic of yeast cell cycle, annotated with cell cycle phase **(B)** Principal component plot of observed cells data colored by assigned cell cycle time, (left) and in-CAHOOTTS predicted trajectories (right) colored by trajectory time, showing model captures cell cycle trajectories. **(C)** Model performance: *R*^2^ and AUPR values for cell cycle predictions with prior, shuffled prior, and no prior conditions across cross-validations. **(D)** Cell cycle gene expression predictions: Expression trajectories for phase-specific genes *HSP150* (M/G1), *PSA1* (G1), *HTA2* (S), and *CIS3* (G2). Gray dots are individual cells from data held out for validation, blue lines show validation trajectories, and red lines show predicted trajectories, starting from *t*_*cc*_ = 0. **(E)** Expression trajectories predicted from an initial state of *t*_*cc*_ = 0 for phase-specific genes *HSP150* (M/G1, blue), *PSA1* (G1, red), *HTA2* (S, green), and *SML1* (G2, orange). Predicted trajectories are shown for the first cycle, where the model has seen training data, and for an additional 632 minutes, totaling 12 hours (8+ cell cycles). **(F)** Oscillation stability analysis: Peak-to-peak distance analysis shows stable oscillatory behavior over extended time periods.

We train a new model using cells collected during the 20 minutes of exponential growth prior to rapamycin treatment. These cells were computationally ordered into 88-minute cell cycle trajectories based on their gene expression profiles (10), reconstructing cell cycle dynamics from cells captured at different cycle stages during the sampling period. The model was provided an initial condition at *t*_*cc*_ = 0 and trained on cell cycle trajectories from *t*_*cc*_ = 0 to *t*_*cc*_ = 88. The cell cycle consists of 4 phases; G1, S, G2, and M-phase, which have different lengths and distinct gene expression patterns (45). The mitotic M-phase results in asymmetric division into two cells, both of which will be in G1-phase; one is a ‘mother’ cell considered continuous to the pre-division cell, and one of which is a new ‘daughter’ cell. The trained in-CAHOOTTS predicts gene expression as a cell goes through multiple cell cycle and mitotic division rounds. This allowed us to evaluate long-term predictions of oscillatory behavior in dynamic equilibrium, extending far beyond training times.

The trained model accurately reconstructs the transcriptome trajectory, starting from *t*_*cc*_ = 0, of the progression through the full cell cycle (Figure 5B). The second and fourth principal components show a characteristic circular pattern that represents progression through the cell cycle (Figure 5B; left). The predicted trajectories PCA plot (Figure 5B; right) recreates the same periodicity and phase relationships as the actual data, demonstrating that the model has learned the underlying dynamics of cell cycle gene expression. Model performance across the transcriptome is quantified by GRN reconstruction AUPR (Figure 5C; top) and by *R*^2^ (Figure 5C; bottom).

Unlike the rapamycin response which follows a monotonic trajectory through transcriptional space, the cell cycle creates a closed loop, requiring continuous oscillatory predictions. We see that model is still able to predict expression profiles that are highly nonlinear across the main phases of the cell cycle (Figure 5D) for phase-specific genes *HSP150* (G1/M), *PSA1* (G1), *HTA2* (S) and *CIS3* (G2). This includes genes which cycle between high expression and low expression (*HSP150, HTA2*) and genes that have spikes followed by slow decrease in expression (*PSA1, CIS3*)

With the goal of making long-term future predictions with these dynamic models, we wanted to see how far into the future this model could predict. We predicted gene expression for 12 hours, encompassing roughly 8 complete cell cycles. Remarkably, the model predicts stable oscillations of gene expression across all phases of the cell cycle (Figure 5E) with consistent periodicity throughout the entire timeframe (Figure 5F). The predicted peak-to-peak distance of 90.90 minutes closely matches the actual cell cycle length of 88 minutes, demonstrating the model has learned the intrinsic timing of cell division. While the oscillatory patterns are stable, the predicted expression magnitudes do dampen compared to early predictions. This likely reflects the model’s tendency toward stable attractors in the learned dynamical system, as the oscillations persist but settle into a more constrained amplitude range over extended time periods.

To determine how far into the future we could predict before the oscillations break down, we extended predictions to 48 hours from *t*_*cc*_ = 0 (Figure 6).The model predictions maintain stable oscillations and also a precise biological periodicity of 87.6 minutes, with phase relationships between genes remaining coordinated across dozens of cell cycles. This demonstrates a key advantage of neural ODEs for biological modeling: unlike many approaches that either blow up exponentially or decay to static steady states, our model maintains meaningful oscillatory dynamics for 30 times longer than training data. This stable limit cycle behavior reflects the true nature of cell cycle dynamics and highlights the underlying mathematical biological mechanisms captured by our framework.

**Figure 6:**
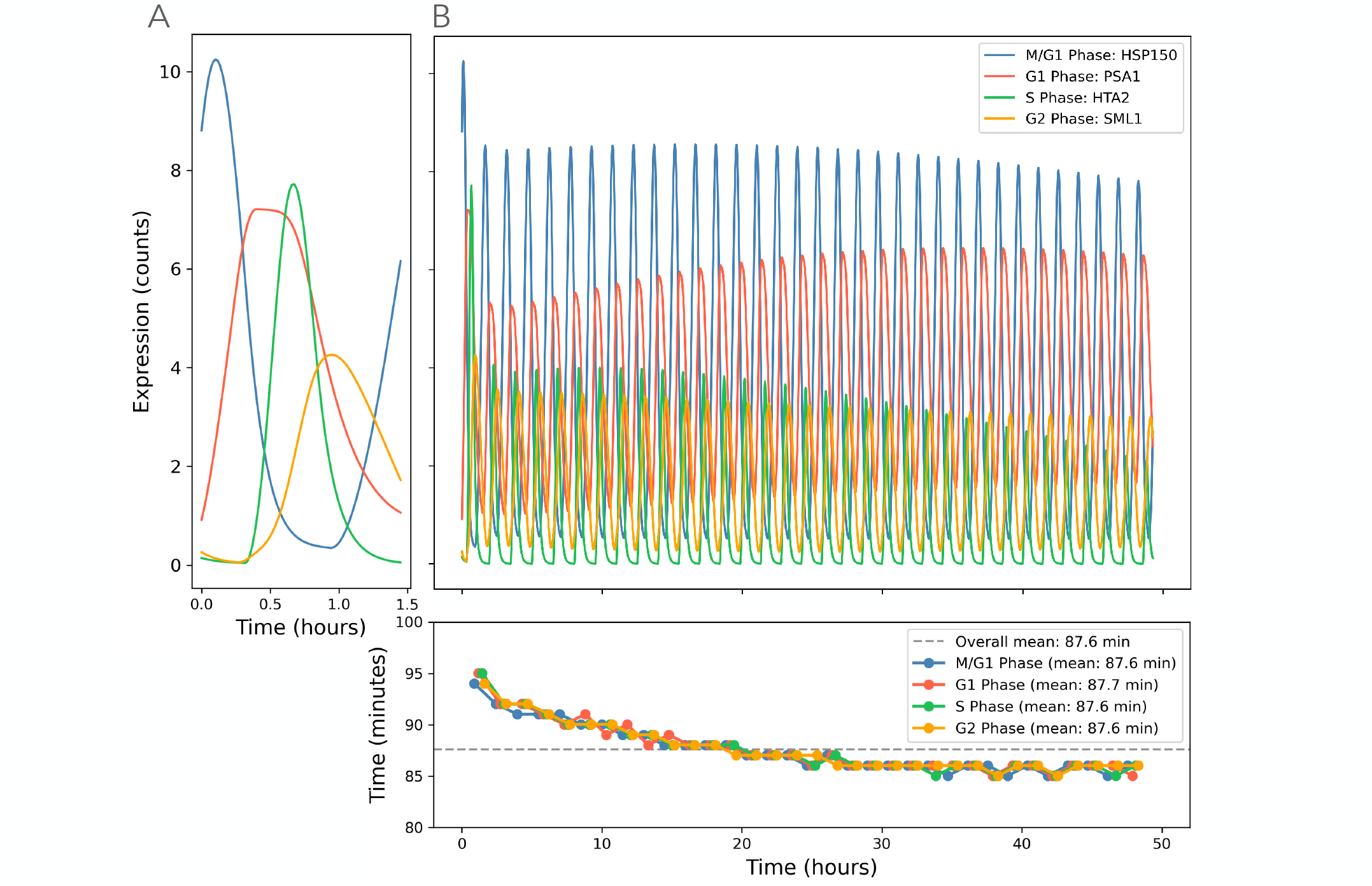
in-CAHOOTTS maintains stable cell cycle oscillations for 48 hours with remarkable biological fidelity. A: Single cell cycle showing phase-specific gene expression patterns for M/G1 (HSP150), G1 (PSA1), S (HTA2), and G2 (SML1) phases during training time of the cell cycle, 87 minutes.B: Long-term predictions extending 48 hours (30× beyond training data) demonstrate sustained oscillatory dynamics with consistent 87.6-minute periodicity. Genes across distinct cell cycle phases show coordinated transcriptional programs maintained across dozens of cell cycles. C: Peak-to-peak distance analysis confirms stable periodicity over the entire 48-hour timeframe, with mean cycle length (87.6 minutes) matching biological cell cycle duration. The model captures true dynamical attractors rather than merely fitting training data, enabling unprecedented long-term biological predictions.

To uncover the gene regulatory network driving cell cycle dynamics, we evaluated the model’s ability to learn biologically meaningful TF-gene interactions. As with the rapamycin analysis, the model was initially trained for 500 epochs without prior knowledge, allowing for purely data-driven discovery of regulatory connections. However, without biological constraints, the model fails to recover known cell cycle regulatory interactions (Figure 5C).

When trained with the prior knowledge adjacency matrix constraining the transcriptional encoder to documented TF-gene interactions, the model successfully recovers cell cycle regulatory networks at a rate 1.6-fold higher than the unconstrained version. Importantly, this biological regularization maintains strong predictive performance for oscillatory dynamics (*R*^2^ values in Figure 5C). The prior-guided model significantly outperforms both the unconstrained model and a shuffled prior control, demonstrating that incorporating biological knowledge enables discovery of meaningful regulatory architecture rather than spurious correlations.

We tested the limits of the framework when cell cycle data is more sparsely sampled (Supplemental Figure S7). We subsampled the 88-minute cell cycle trajectories at intervals of 11 and 4 minutes and compared performance against the original 1-minute sampling method. R2 values remain relatively high across these sampling densities when using the prior, suggesting the model can capture oscillatory dynamics even with reduced temporal resolution. However, the model fails to converge when trained with intervals beyond 11 minutes, indicating that the complex oscillatory dynamics of the cell cycle require sufficient temporal sampling to be accurately learned by in-CAHOOTTS.

Notably, the cell cycle regulatory network shows distinct patterns from the rapamycin response network, reflecting the different transcriptional programs governing oscillatory cell division versus stress response. The predicted biophysical parameter estimates (Supplemental Figure S14) and transcription factor activities (Supplemental Figure S16) exhibit clear oscillatory behavior consistent with the cyclical nature of cell division. Phase-specific TF activity patterns align with known cell cycle biology, demonstrating the model’s ability to learn context-specific regulatory mechanisms underlying different biological processes.

## 5 Discussion

We have demonstrated that biophysically-motivated neural ODEs can successfully model complex biological dynamics while maintaining interpretability. We aimed to create a framework that could not only capture cellular responses to perturbations but decompose them into interpretable biophysical processes, uncover the underlying gene regulatory networks, and extend predictions far beyond training data across diverse biological contexts while maintaining full biological interpretability. Our framework represents a significant advance by combining accurate prediction with mechanistic understanding, a critical requirement for studying living systems where we must understand not just what happens, but why. The success of in-CAHOOTTS across multiple biological contexts, rapamycin stress response and cell cycle dynamics, demonstrates the fundamental utility of neural ODEs for biological modeling. These models generalize across systems because they capture the underlying mathematical structure of biological processes: differential equations governing change over time. The ability to achieve stable predictions 30 times longer than training data while maintaining biological realism further supports the broad potential of these approaches in systems biology.

A key advantage of our neural ODE approach is that it implicitly learns both RNA velocity and mRNA decay rates directly from gene expression data, without requiring precomputed estimates of these biophysical parameters. In contrast, the previous RNN-based architecture developed for the time-series data used in this study, (10), required pre-estimated RNA velocity and mRNA decay rates as inputs during training. By learning these parameters end-to-end from the expression trajectories, our model not only outperforms the RNN approach in AUPR and long-term predictive capabilities, it also provides a more unified framework that discovers the underlying biophysical dynamics without relying on potentially noisy or method-dependent pre-processing steps. This explicit implementation demonstrates the power of neural ODEs to capture the continuous dynamics of gene regulatory systems while simultaneously inferring the mechanistic parameters that drive observed expression changes.

Our approach of training neural ODEs to predict from single time points, rather than embedding multiple time points in the forward pass, pushes the model toward its true predictive limits. This design choice forces the model to learn the vector field structure of the dynamical system, enabling robust extrapolation. However, this comes with challenges, neural ODEs can be highly resistant to regularization, and incorporating biological priors here required novel soft constraint approaches rather than traditional penalization methods.

We demonstrate that interpretable deep learning is not just preferable but essential for biological applications. The disconnect we observed between transcription factor activity and expression, with dramatic activity changes occurring despite stable mRNA levels, would be invisible to black-box approaches. This biological insight emerged directly from our interpretable architecture and represents the kind of mechanistic understanding necessary for advancing biological knowledge. By successfully integrating prior biological knowledge with flexible machine learning architectures, we show that the most powerful biological models emerge when mechanistic understanding guides statistical learning (16; 18; 28).

A fundamental limitation of our current approach is the deterministic nature of ODEs, while biology is inherently stochastic. This limitation highlights a broader challenge in dynamic biophysical modeling: how to incorporate biological stochasticity as a feature containing crucial mechanistic information about underlying biophysical processes. Stochasticity is not merely noise to be filtered out, but rather the mechanism by which biological systems explore state space, make decisions, and maintain function (58; 16; 18; 59). Future work should explore Neural Stochastic Differential Equations (Neural SDEs) to model transcriptional bursting, cellular noise, and stochastic switching events. This could enable modeling of cellular bifurcations in development and perturbation responses while maintaining the interpretable structure we’ve established. Such an approach would align with generating function frameworks that model biological stochasticity and technical noise in a unified fashion, providing a more complete picture of cellular dynamics (16; 18; 33; 59). Ultimately, this would move us toward models that capture how living cells exploit variability as a fundamental feature rather than a limitation to overcome.

Our method currently requires relatively dense, high-quality time series data, a rarity in single-cell biology. Generative neural ODEs could address this limitation by learning continuous representations without predefined trajectories, potentially enabling application to sparser datasets. Additionally, moving beyond trajectory means to full generative modeling could capture cell-to-cell variability more accurately (60; 61; 35; 34; 18; 25). Importantly, we need to test this framework on larger, more complex systems beyond *S*. cerevisiae to evaluate scalability. The question remains whether our approach can maintain interpretability and biological realism when applied to mammalian systems with orders of magnitude more genes and regulatory complexity (12; 62; 63).

This work represents a step toward AI that not only predicts biological outcomes but explains the mechanisms driving them. As we demonstrated, the greatest challenge was not building accurate models, but ensuring they learned biologically meaningful representations. The successes of in-CAHOOTTS exemplify a broader paradigm shift in computational biology toward mechanistic-statistical fusion. Like the PHOENIX framework and biVI approaches described in recent literature, our method demonstrates that the most powerful biological models emerge when mechanistic domain knowledge guides flexible statistical learning architectures (28; 18). This mechanistic models represent a departure from purely black-box machine learning approaches that, while predictive, fail to provide the mechanistic insights essential for biological understanding (39; 19). This trend toward interpretable, mechanism-aware AI is exemplified by recent developments like the AIDO foundation models, which demonstrate how large-scale biological modeling can maintain mechanistic grounding while achieving broad applicability (64). Such approaches suggest a future where AI and domain expertise are deeply integrated, with models serving as hypothesis-generating tools that enhance rather than replace biological understanding.

## 6 Methods

### 6.1 Data Preprocessing and Trajectory Construction

#### 6.1.1 Dataset

Single-cell RNA sequencing data was obtained from GEO record GSE242556 for *Saccharomyces cerevisiae* cells responding to rapamycin treatment (10). The dataset contains 173,361 cells continuously sampled over 80 minutes, with 60 minutes following rapamycin treatment, and additional cell-cycle time axis spanning 88-minute cycles, across 5,726 detected genes. Cells are annotated with inferred time since rapamycin treatment (*t*) and inferred time in the cell cycle (*t*_*cc*_) Bulk RNA sequencing data for rapamycin response was obtained from GEO record GSE242556.

#### 6.1.2 Data Normalization and Scaling

Integer transcript counts were normalized per cell to the median count depth (3,099 reads per cell) in scanpy (65). Normalized expression values were scaled using a modified RobustScaler (Equation 15 from (10)). Time axes were linearly scaled to the range [0,1] to standardize temporal inputs across different experimental conditions and unscaled post training.

#### 6.1.3 Rapamycin Model Training

##### Training trajectories

During pretraining, the model received 10 cells from minutes 0-9 as initial conditions, with 30 additional randomly selected time points between 10-60 minutes serving as regularization targets. For prior-guided training, cells were arranged into 60-minute trajectories with one-minute sampling intervals. Due to high expression variance across timepoints, 20 trajectories were averaged together to create a single lower-noise trajectory for neural ODE learning.

##### Testing trajectories

Validation data consisted of single 60-minute trajectories with cells sampled at one-minute intervals.

##### Trajectory resampling

To prevent overfitting, trajectories were resampled after each training epoch.

#### 6.1.4 Cell Cycle Model Training

##### Training trajectories

Given the 88-minute cell cycle length (*t*_*cc*_), the model was trained on three sequential 29-minute trajectory segments (0-29, 29-58, and 58-87 minutes) to reduce learning complexity. The same two-stage training protocol was applied: 500 epochs without priors followed by 600 epochs with prior regularization. During pretraining, data was resampled twice per epoch to increase data exposure.

##### Validation trajectories

Post-training validation used continuous 88-minute trajectories starting from *t*_*cc*_ = 0.

#### 6.1.5 Sparse Sampling Experiments

The framework was retrained to test if the neural ODE framework could still be predictive on less continuous data. The full trajectories with cells sampled 1-minute apart were subsampled across different intervals: 30, 20, 10, 5, and 1 minute for the rapamycin and 11, 4, and 1 minute for the cell cycle.

Sparse temporal sampling was performed at nominally regular intervals of *s* minutes, with uniform random jitter *U* (−*s/*4, *s/*4) applied to all sampling points except the first to prevent systematic bias.

#### 6.1.6 Prior Knowledge Network

A prior knowledge network *P* (*P* ∈ 0, 1) consists of 14,018 regulatory edges, connecting 158 regulatory TFs to 5,726 yeast genes was obtained from the YEASTRACT database (53). For testing purposes, we use an existing gold standard regulatory network of high-confidence interactions, which are a subset of the YEASTRACT prior network (29).

#### 6.1.7 Model Architecture Details

The in-CAHOOTTS framework consists of two parallel neural networks with the following specifications:

**Transcription Model (***α*): input layer: 5,726 genes, hidden layers: 158 → 158 → 158 (3 hidden layers, corresponding to TF dimensionality), output layer: 5,726 genes (transcription rates). Activation functions: GeLU for hidden layers, ReLU for output layer (50; 51). The latent dimensionality of 158 corresponds to the number of transcription factors in the prior knowledge network, enabling biological interpretation of the learned representations.

**Decay Model (***λ*): Input layer: 5,726 genes, Hidden layers: 158 → 158 (2 hidden layers), Output layer: 5,726 genes (decay rates), Activation functions: Softplus throughout, ReLU for output layer.

**Vanilla Neural ODE** Input layer: 5,726 genes, Hidden layers: 158 → 158 (4 hidden layers), Output layer: 5,726 genes (decay rates), Activation functions: Softplus throughout, ReLU for output layer.

### 6.2 Model Training

The in-CAHOOTTS model was trained to minimize mean squared error (MSE) between predicted and actual gene expression values:

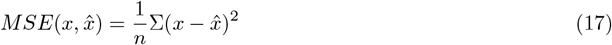

for all trajectories from 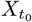 to 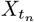. During the forward pass, 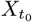 is linearly embedded into both the transcription and decay models, while gradients during backpropagation update model weights to minimize the error between 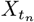 and 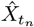.

#### 6.2.1 Two-Stage Training Protocol

##### Stage 1: Data-driven pretraining (500 epochs)

The model was initially pretrained for 500 epochs without biological priors, optimizing purely on gene expression prediction. This data-driven approach allows the neural ODE to learn cellular dynamics directly from single-cell expression data. For regularization, the model was trained on the first 10 minutes of data (sampled at one-minute intervals from 0-9 minutes) plus 30 additional randomly selected time points between 10-60 minutes, rather than using full 60-minute trajectories sampled at one-minute intervals. Following pretraining, the weights of the decay model (*λ*) were frozen to preserve the estimated mRNA decay rates.

##### Stage 2: Prior-guided training (600 epochs)

After pretraining, biological prior knowledge was incorporated into the transcriptional model and trained for an additional 600 epochs. The prior adjacency matrix guided the initialization of first-layer encoder weights in the transcriptional model, with scale normalization applied to match the distribution of randomly initialized weights. This embeds the data into a latent space with dimensionality equal to the number of transcription factors.

#### 6.2.2 Prior Knowledge Regularization

During prior-guided training, we apply a regularization term that penalizes deviation from documented biological interactions:

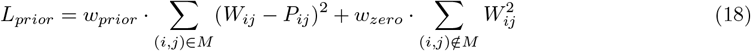

where *W*_*ij*_ are the weights of the first encoder layer, *P*_*ij*_ are the prior knowledge values from the adjacency matrix, *M* is the mask of non-zero entries in the prior matrix, and *w*_*prior*_ and *w*_*zero*_ are regularization coefficients. This approach encourages the model to preserve documented TF-gene interactions while allowing data-driven discovery of novel regulatory connections.

### 6.3 Hyperparameter Optimization

Model parameters were optimized to maximize *R*^2^ values for predicted gene expression, with 20% of cells withheld for validation and 80% used for training. The model was simultaneously tuned for optimal Area Under the Precision-Recall Curve (AUPR) performance, with 20% of known regulatory connections withheld from training and used for validation.

#### 6.3.1 Regularization

- **Transcriptional model:** 20% dropout rate applied to the encoder layer, 10% dropout rate applied to the latent layer
- **Learning rate:** *γ* ∈ {10^−3^, 5 *×* 10^−3^, 10^−4^, 5 *×* 10^−5^, 10^−5^}
- **Weight decay:** *λ* ∈ {10^−3^, 10^−4^, 10^−5^, 10^−6^}

Hyperparameters were tuned to optimize validation AUPR and *R*^2^ simultaneously.

### 6.4 Implementation Details

The model was implemented in PyTorch using the torchdiffeq package for ODE solving. Training was performed on CUDA-enabled GPUs with gradient checkpointing for memory-efficient processing of larger datasets.

#### 6.4.1 ODE Solver Configuration

- **Solver method:** 4th-order Runge-Kutta with adjoint method for efficient gradient computation
- **Optimizer:** Adam with learning rate scheduling (reduction factor 0.9 when validation loss plateaus, patience=30)
- **Cross-validation:** 5-fold cross-validation was conducted for all experiments

### 6.5 Evaluation Metrics

#### 6.5.1 Expression Prediction Performance

*R*^2^ **(Coefficient of Determination):** Measures the proportion of variance in observed gene expression explained by model predictions:

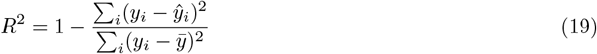

#### 6.5.2 Gene Regulatory Network Recovery

##### Explained Relative Variance (ERV)

Trained models are interpreted to provide evidence for biological TF-to-gene regulatory relationships. For each transcription factor, a TF-reduced model is created by removing that TF from the full model. ERV is calculated as one minus the ratio of the RSS of the TF-reduced model to the RSS of the full model:

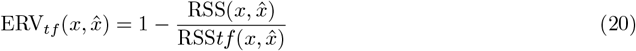

This procedure is repeated for each TF to calculate a genes × TFs ERV matrix (ERV ∈ *R*^*g×k*^ : ERV ≤ 1), where *g* is the number of genes and *k* is the number of transcription factors.

##### AUPR (Area Under Precision-Recall Curve)

TF-to-gene regulatory edges are ranked by explained relative variance and compared to known TF-to-gene regulatory relationships derived from literature. Causal network inference performance is evaluated as Area Under the Precision-Recall curve (AUPR), which accounts for the highly imbalanced nature of regulatory network data.

#### 6.5.3 Visualizations

All visualizations were generated in matploblib (66).

## 6.6 Acknowledgements

We thank the members of the Bonneau Lab for insightful discussions and feedback on this manuscript. We also thank the staff of the NYU IT High Performance Computing and Flatiron Institute Scientific Computing Core. We are additionally grateful to Ji Won Park and Daniel Berenberg for insightful discussions related to this work.

## 6.7 Code Availability

Code for the in-CAHOOTTS framework and all analyses is available at https://github.com/maggiealetha/incahootts.

## 6.9 Author Contributions

Conceptualization: MBA, CAJ, RB, DG; Methodology: MBA, CAJ; Software: MBA, CAJ; Formal analysis: MBA; Investigation: MBA, CAJ; Validation: MBA; Resources: CAJ; Data curation: MBA, CAJ; Visualization: MBA; Writing – original draft preparation MBA, CAJ, RB, DG; Writing – review and editing: MBA, CAJ, RB, DG; Funding acquisition: RB, DG; Project administration: MBA, CAJ, RB, DG.

---

**Supplemental Figure S1:**
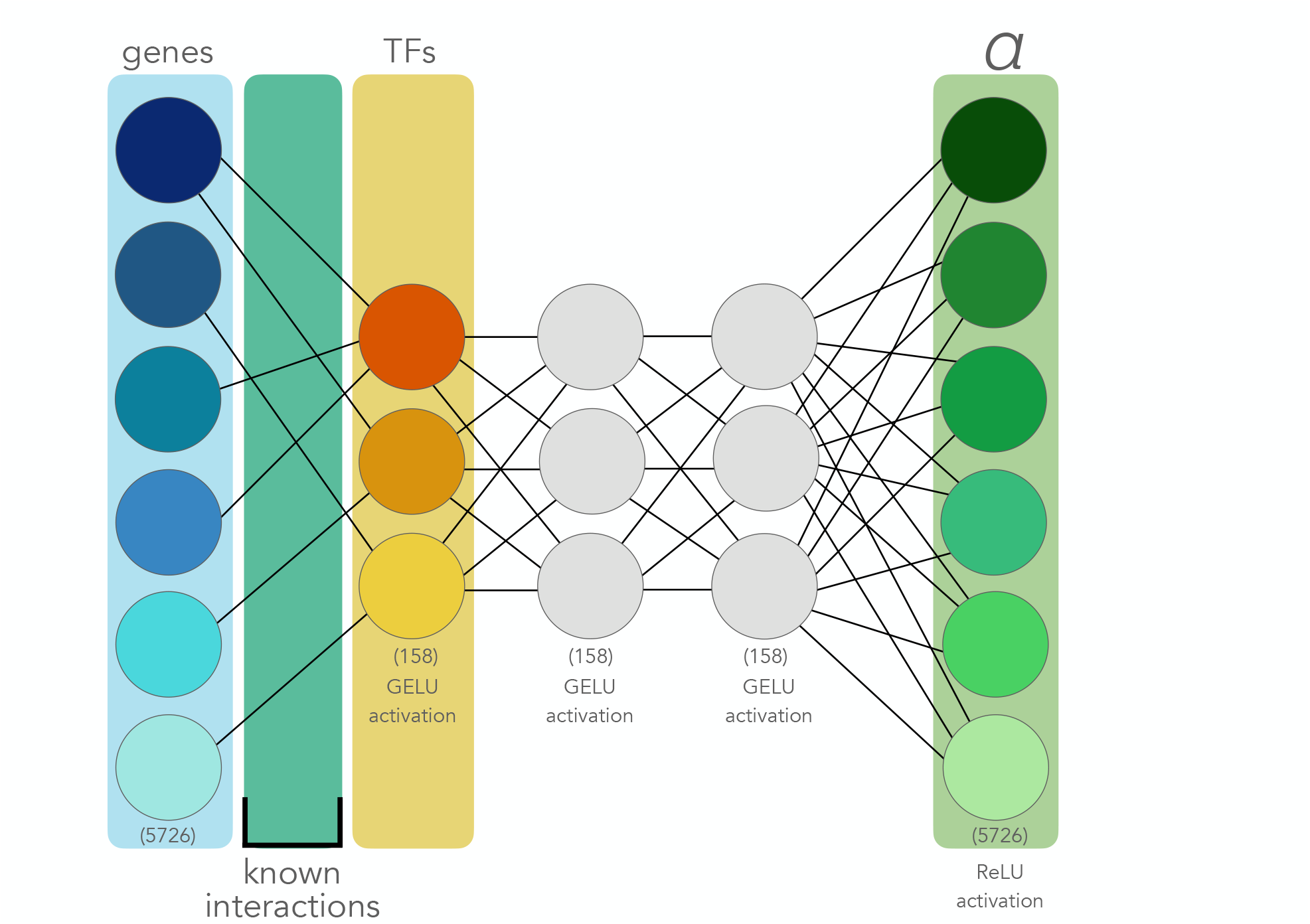
Transcriptional Model Architecture. Schematic of the interpretable deep learning architecture for estimating mRNA transcription rates (*α*). Gene expression (left, blue, dimensionality 5726) is encoded through known TF-gene interactions (teal) into a latent space interpretable as transcription factor activities (yellow, TFs, dimensionality 158 for all TFs) with GELU activations applied both after encoding and between all latent layers. The architecture uses two intermediate hidden layers (gray) before decoding to transcription rate outputs (right, green *α*). A ReLU activation is applied so so the outputs are interpretable as positive mRNA transcription rates. Black lines indicate learned connections, with the encoder weights constrained by prior biological knowledge, in the form of an adjacency matrix, to maintain interpretability of the latent space as transcription factor activities.

**Supplemental Figure S2:**
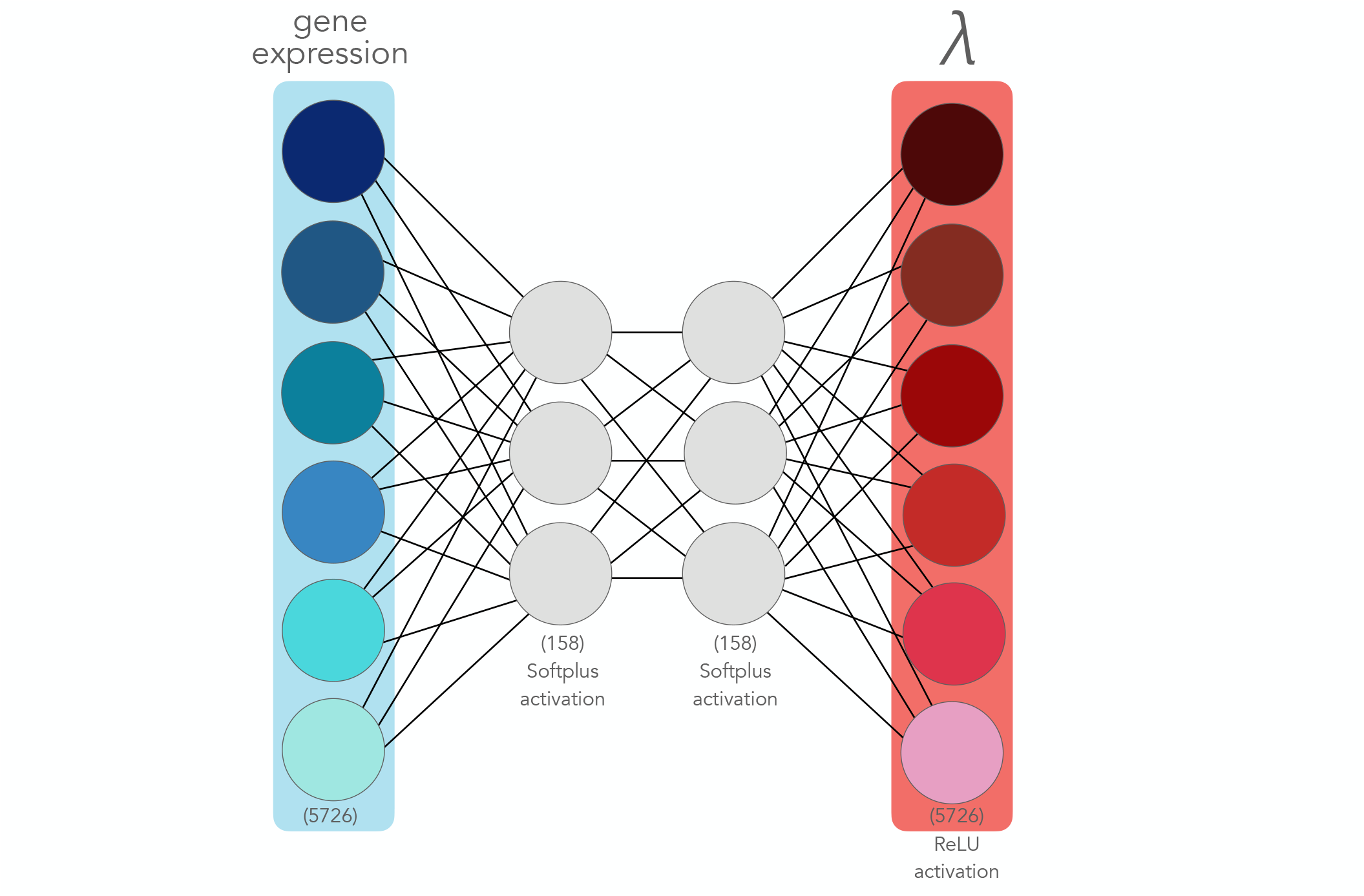
Decay Model Architecture. Schematic of the interpretable deep learning architecture for estimating mRNA decay rates (*λ*) per gene. Gene expression (left, blue, dimensionality 5726) is encoded into a latent space (dimensionality 158) with Softplus activations applied both after encoding and between latent layers. The architecture uses two intermediate hidden layers (gray) before decoding to mRNA decay rate outputs (right, red *λ*). A ReLU activation is applied so the embeddings are interpretable as positive mRNA decay rates. Black lines indicate learned fully connected weights, as this model functions as a black box.

**Supplemental Figure S3:**
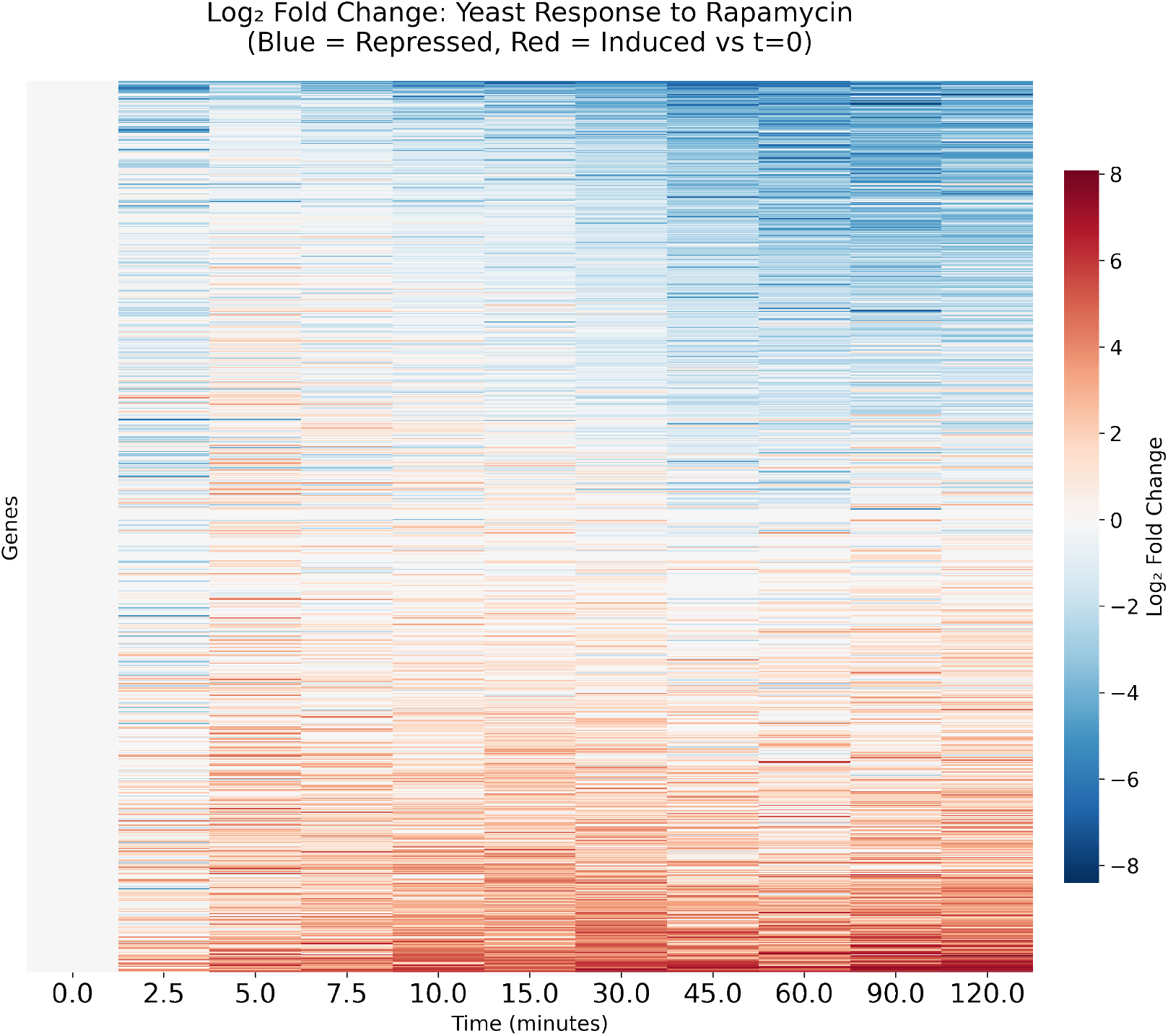
Heatmap of bulk gene expression responding to rapaymcin for 120 minutes. *Log*_2_ fold change of gene expression in S. cerevisiae over 120 minutes following rapamycin treatment (blue = repressed, red = induced relative to t=0). Genes are grouped by functional categories showing coordinated transcriptional responses: ribosomal protein genes (RPL3, etc.) are rapidly downregulated following TOR pathway inhibition, while stress response and metabolic genes (GCN4, GDH1, GAP1, GLN1, MEP2) show upregulation. The heatmap demonstrates the temporal dynamics of the rapamycin response used to train the in-CAHOOTTS model, with clear separation between growth-related genes that are repressed and stress-adaptation genes that are activated.

**Supplemental Figure S4:**
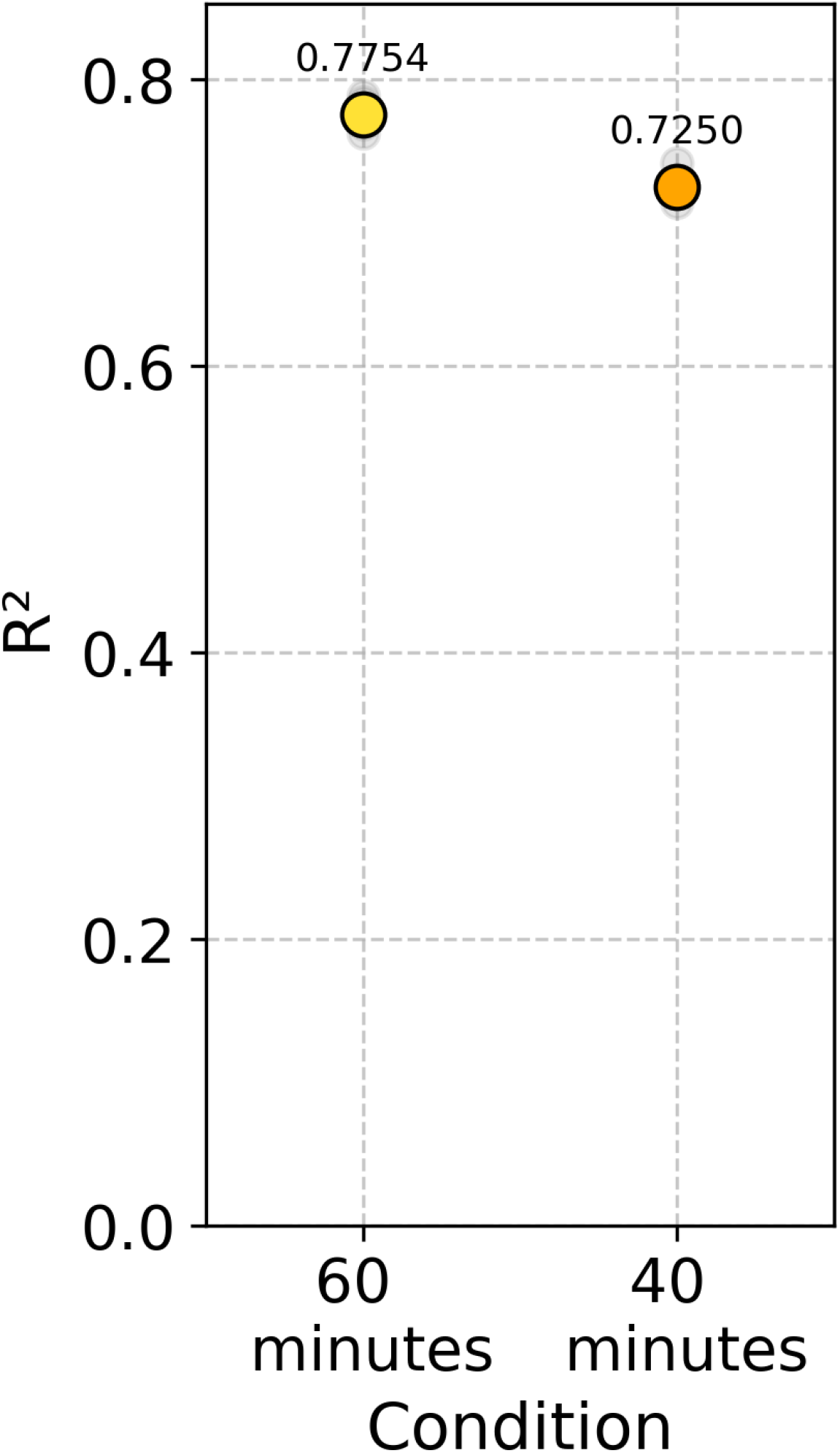
Predictive Performance of Withheld Trajectories. Coefficient of determination (R2) comparison between in-CAHOOTTS trained on 60 minutes of data vs 40 minutes of data (withholding minutes 40-60) on rapamycin response data. in-CAHOOTTS achieves 0.775 R2 compared to 0.725 for the withheld data, demonstrating the model can still achieve high performance on predictions beyond training times.

**Supplemental Figure S5:**
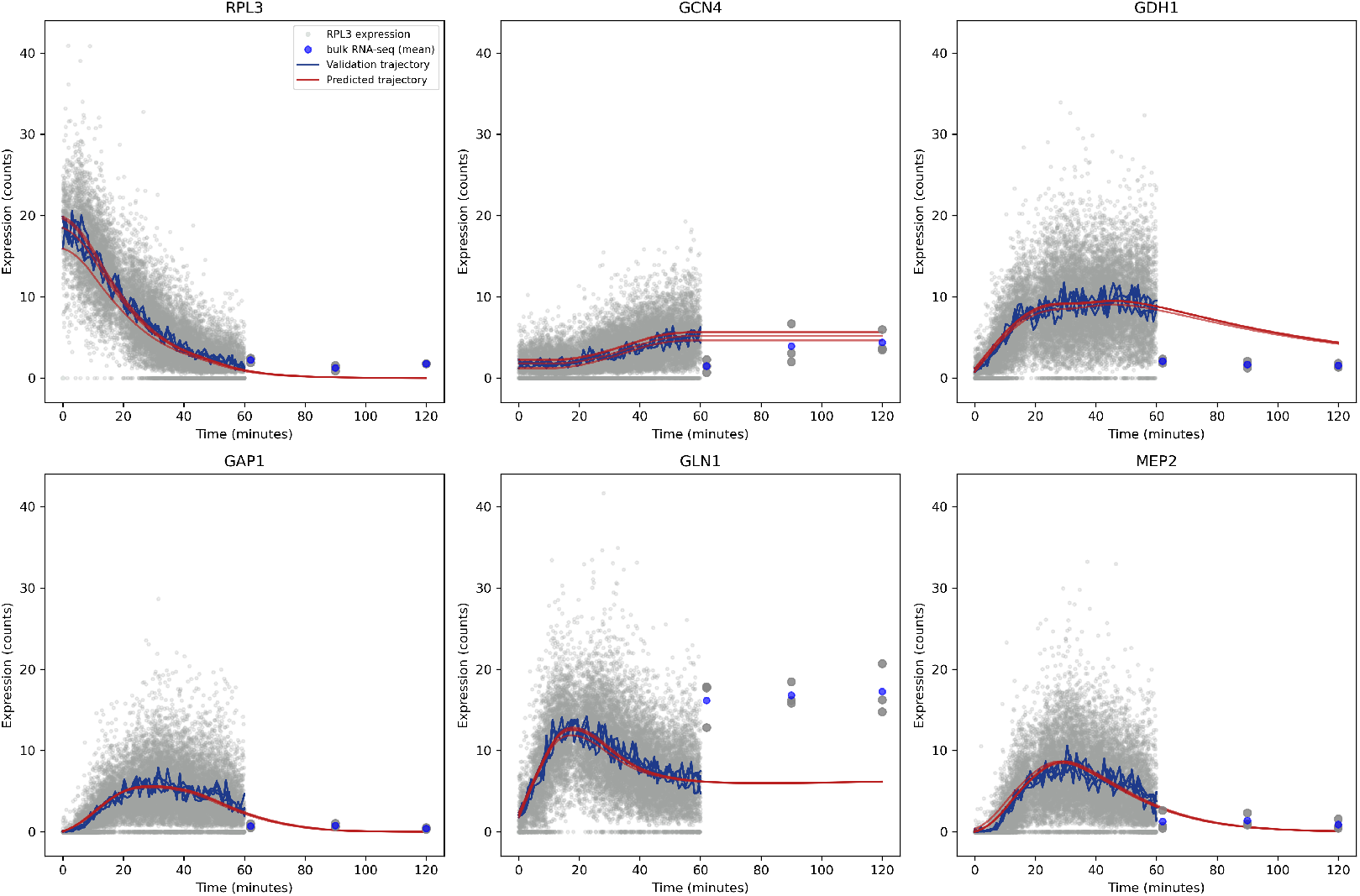
Long-term rapamycin response predictions versus bulk expression data. Comparison of in-CAHOOTTS model predictions (red lines) with bulk RNA-seq data (blue points) for key rapamycin-responsive genes over 120 minutes. Gray dots represent single-cell validation trajectory data used for model training (0-60 minutes). The model successfully captures both the magnitude and temporal dynamics of gene expression changes for ribosomal genes (RPL3), stress response transcription factors (GCN4), amino acid transporters (GAP1, GCN1), and metabolic genes (MEP2). Predictions extend beyond the single-cell training timeframe (60 minutes) and accurately match independent bulk expression measurements, demonstrating the model’s ability to generalize across experimental platforms and predict long-term cellular responses to perturbations.

**Supplemental Figure S6:**
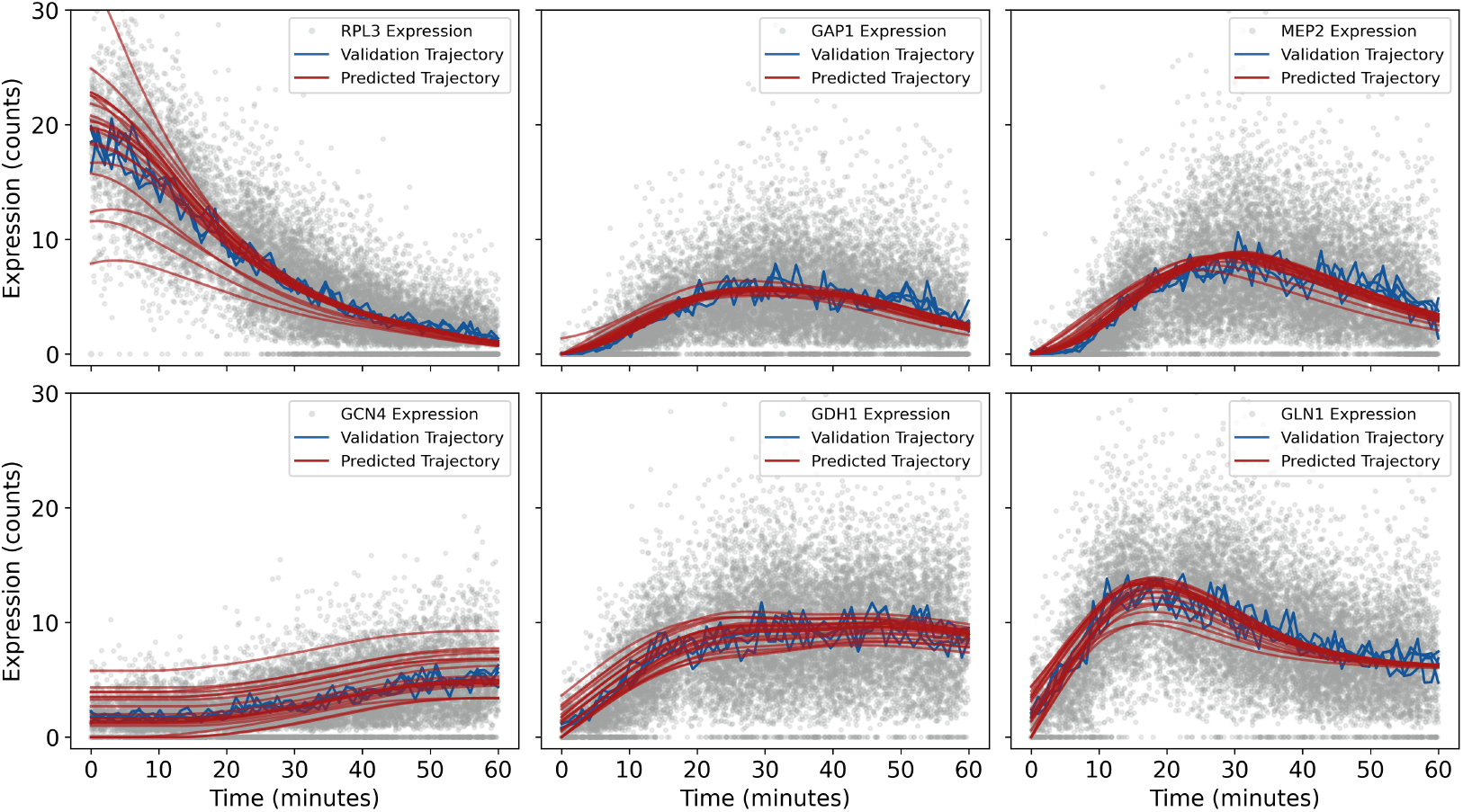
Trajectory Predictions from All Initial Conditions Rapamycin. Predicted trajectories starting the model from initial conditions outside of the mean trajectory space on validation data. Gray dots are individual validation cells (*n* = 19800); blue lines are the average of 20 cells from 1 minute windows, assembled into a validation trajectory (t=0 to t=60 minutes). Red lines are predicted trajectories (t=0 to t=60 minutes at 1 minute intervals).

**Supplemental Figure S7:**
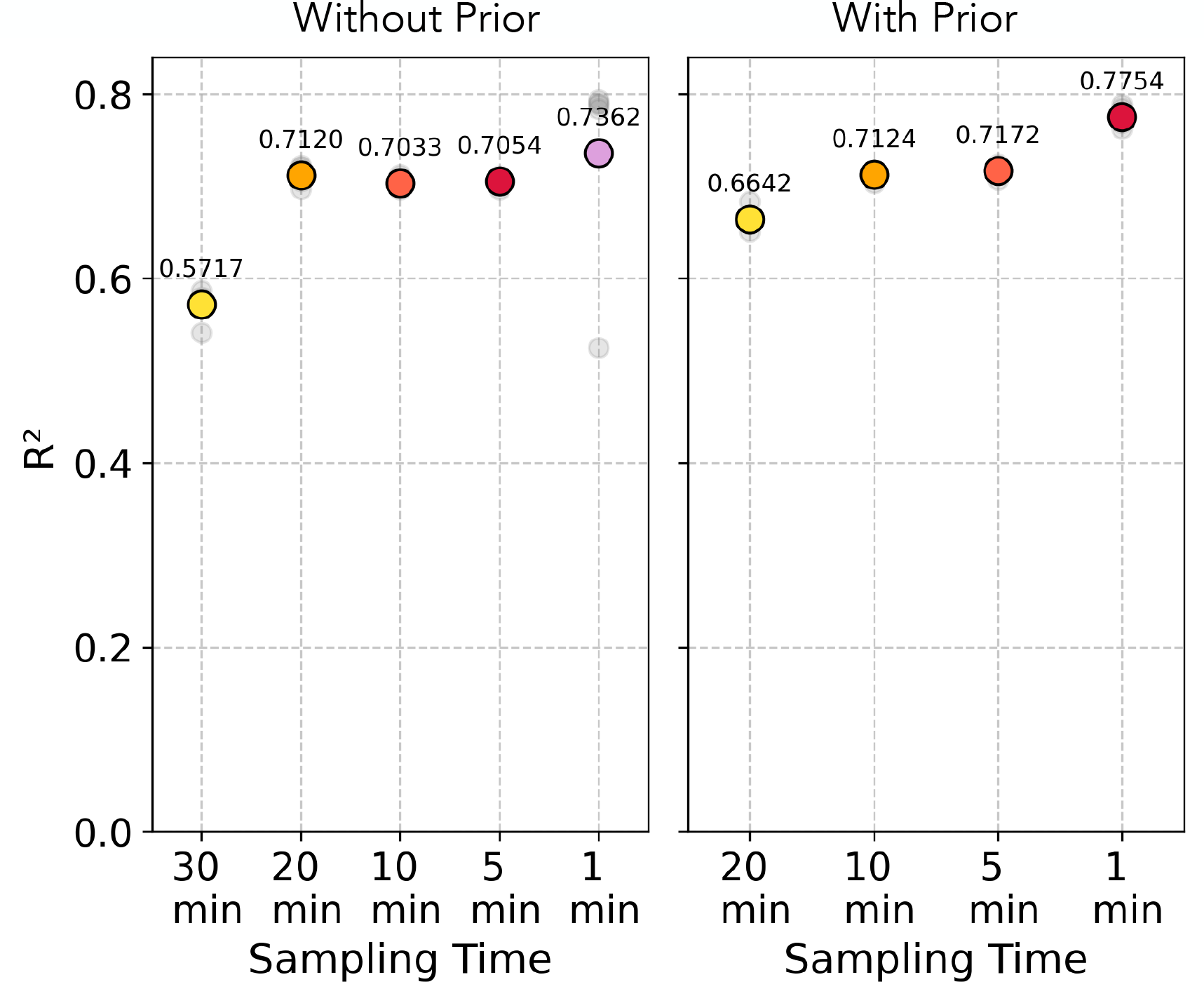
Predictive Capabilities from Sparsely Sampled Trajectories for Rapamycin. Cells are arranged into trajectories of 60 minutes, with cells sampled every 30 minutes, 20 minutes, 10 minutes, 5 minutes and 1 minute apart from the trajectory. A scaling stochastic term (see Methods) based on sampling distance was added around each sample time-bin to avoid overfitting. The neural ODE failed to converge on trajectories sampled sparser than 20 minutes with the prior. However, the R2 values remain high relatively high, suggesting the model can still make predictions on datasets less continuously sampled than ours, and that the model can smoothly interpolate predictions at unseen times.

**Supplemental Figure S8:**
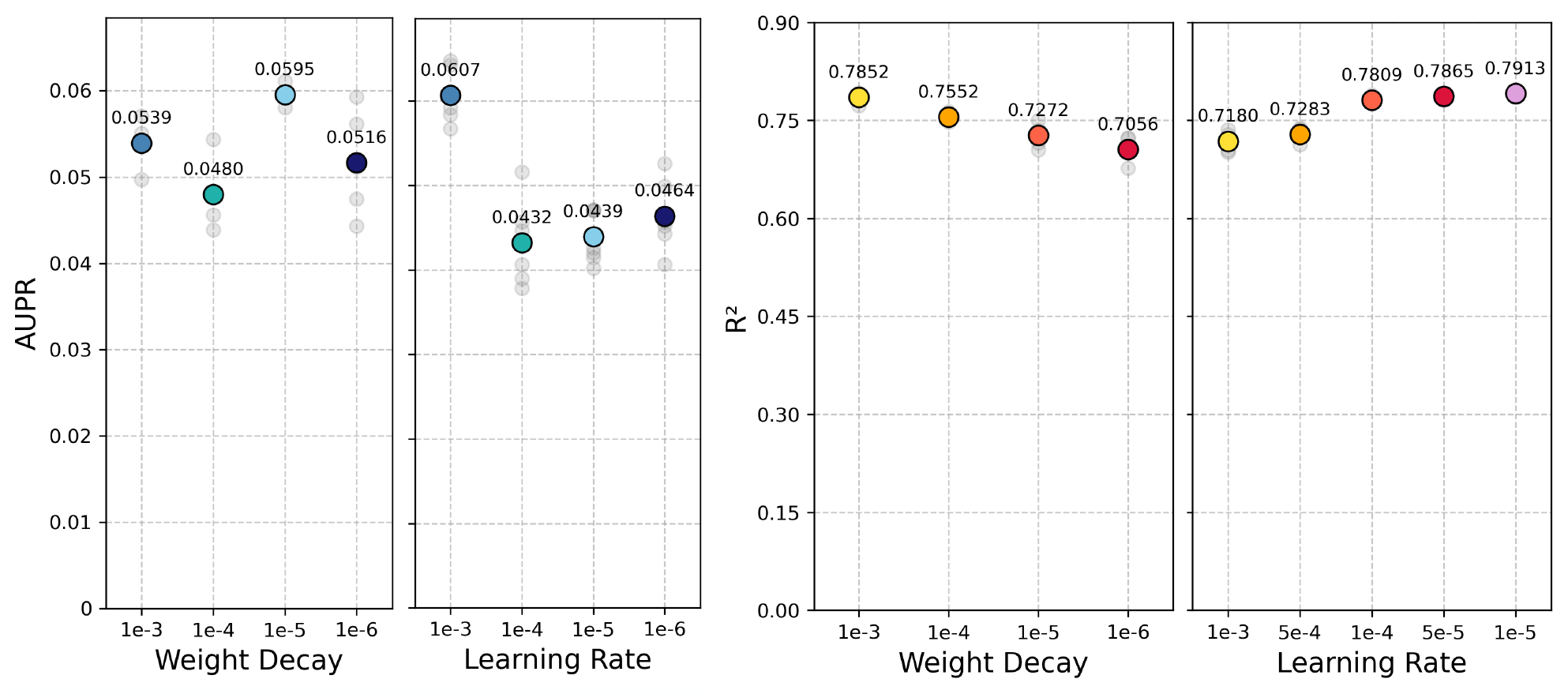
Hyperparameter Grid Search for Rapamycin Response. Individual cells are arranged into trajectories of 60 minutes with cells selected one-minute apart. Performance quantified by AUPR and R2 across 5-cross validations. Performance is shown for a range of Adam optimizer learning rates (*γ*) with weight decay (*λ*) held constant at 1*e*^−^5, and a range of weight decays with learning rate held constant at 1*e*^−^3.

**Supplemental Figure S9:**
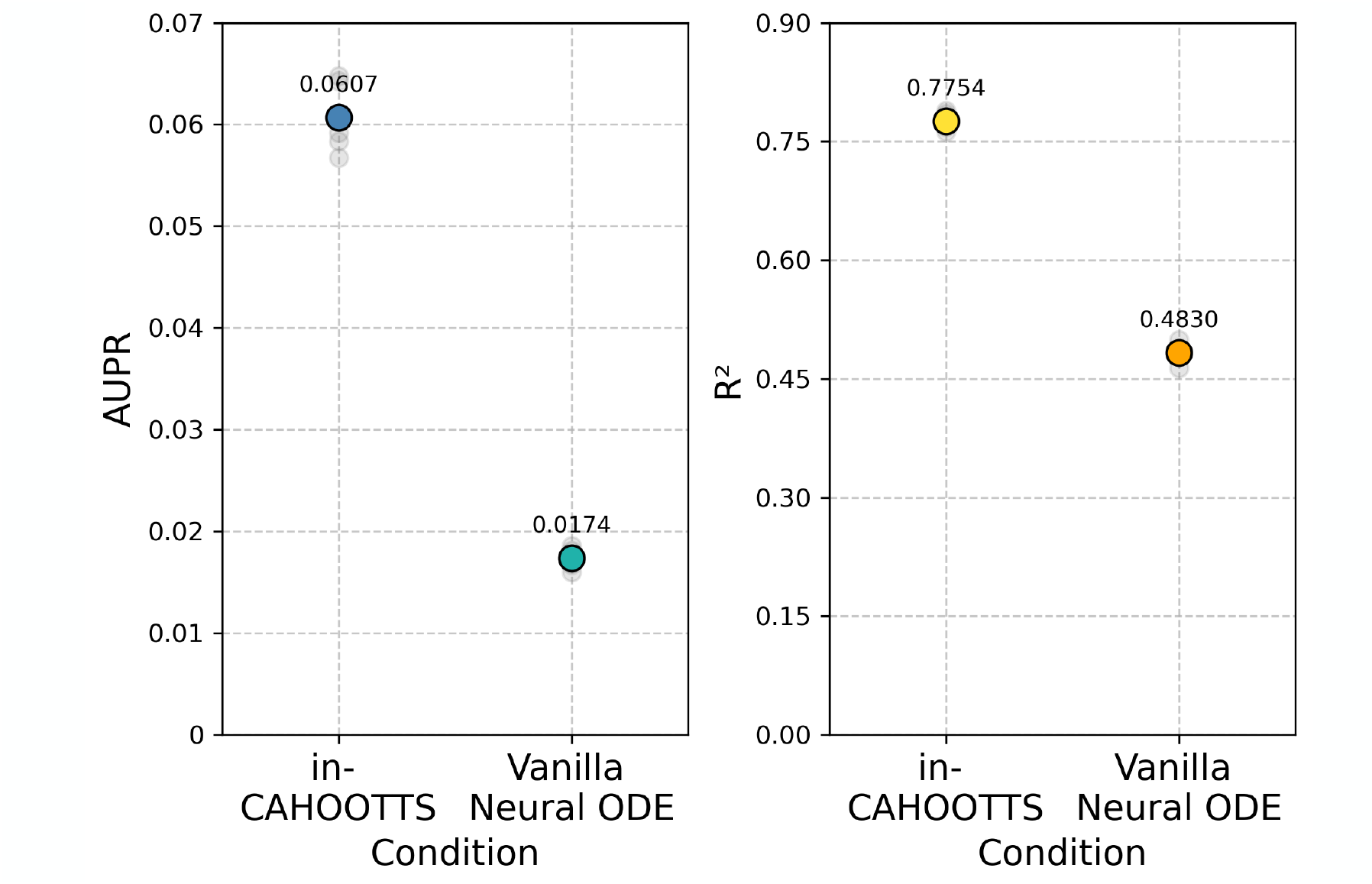
Vanilla Neural ODE vs in-CAHOOTTS. Area Under Precision-Recall Curve (AUPR) comparison for gene regulatory network recovery between in-CAHOOTTS and vanilla Neural ODE. in-CAHOOTTS achieves 0.0607 AUPR compared to 0.0174 for vanilla Neural ODE, representing a 3.5-fold improvement in regulatory network inference. The substantial improvement demonstrates that interpretable architectures with prior biological knowledge are essential for recovering meaningful regulatory relationships rather than spurious correlations. Coefficient of determination (R2) comparison between in-CAHOOTTS and vanilla Neural ODE on rapamycin response data. in-CAHOOTTS achieves 0.775 R2 compared to 0.483 for vanilla Neural ODE, demonstrating a 60 percent improvement in predictive accuracy. The superior performance highlights the importance of biophysical decomposition (*α*/*λ*) and prior biological knowledge integration in capturing gene regulatory dynamics.

**Supplemental Figure S10:**
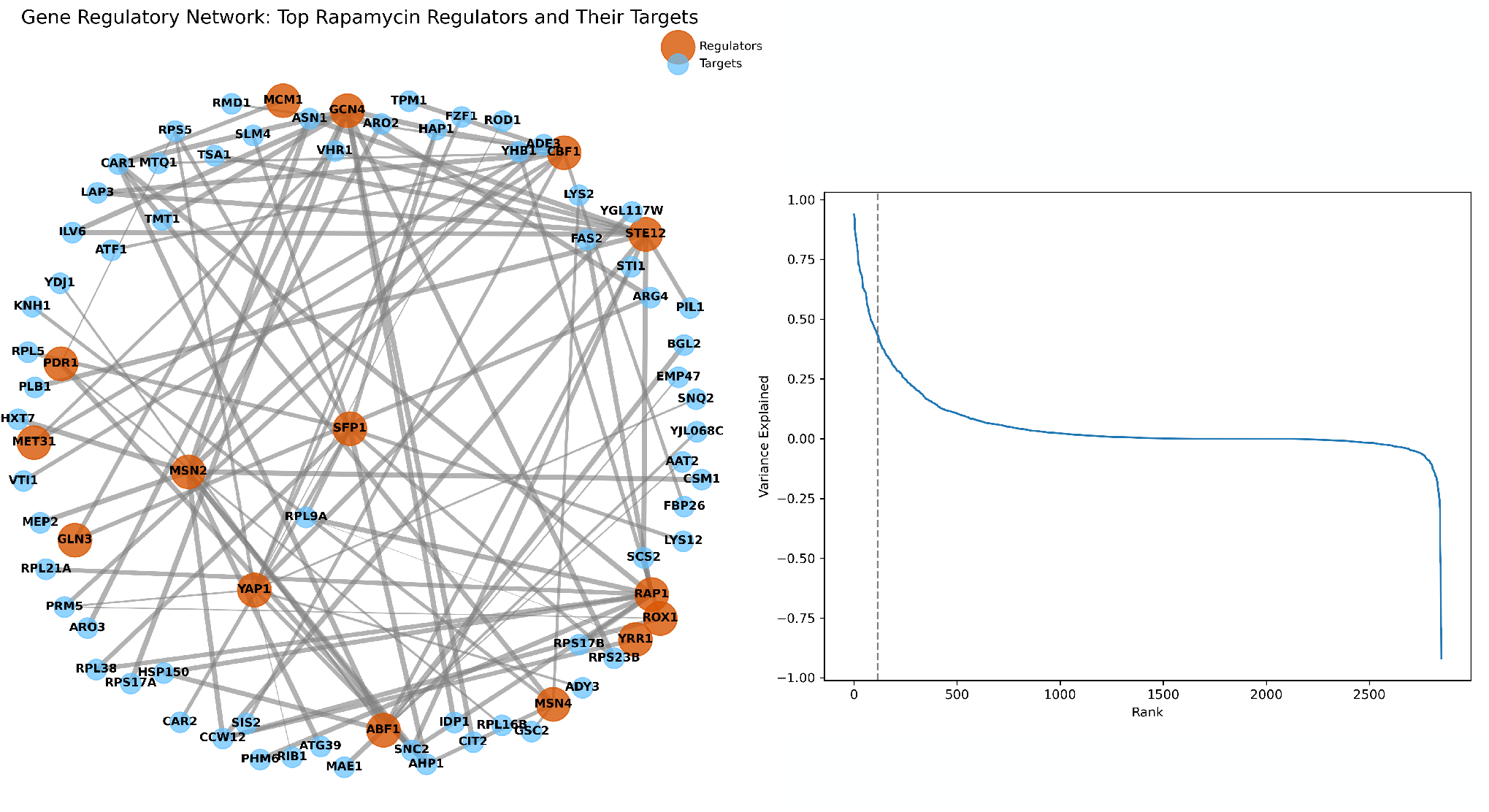
Inferred gene regulatory network reveals key transcription factors driving rapamycin response. Gene regulatory network inferred by in-CAHOOTTS showing the top rapamycin-responsive transcription factors (orange nodes) and their target genes (blue nodes). Network connections represent high-confidence regulatory relationships learned by the model through explained relative variance analysis, constrained to only connections actually existing within the gold standard gene regulatory network. Key hub transcription factors include stress response regulators (MSN2, GCN4), ribosomal biogenesis controllers (ABF1), and metabolic regulators (STE12, CBF1). The inset shows the elbow plot used to determine the optimal number of connections to include in the network visualization, with the elbow at approximately 116 connections capturing the most significant regulatory relationships. Gray edges indicate regulatory interactions constrained by prior biological knowledge, demonstrating how the model recovers known regulatory architecture while discovering context-specific activation patterns during rapamycin treatment.

**Supplemental Figure S11:**
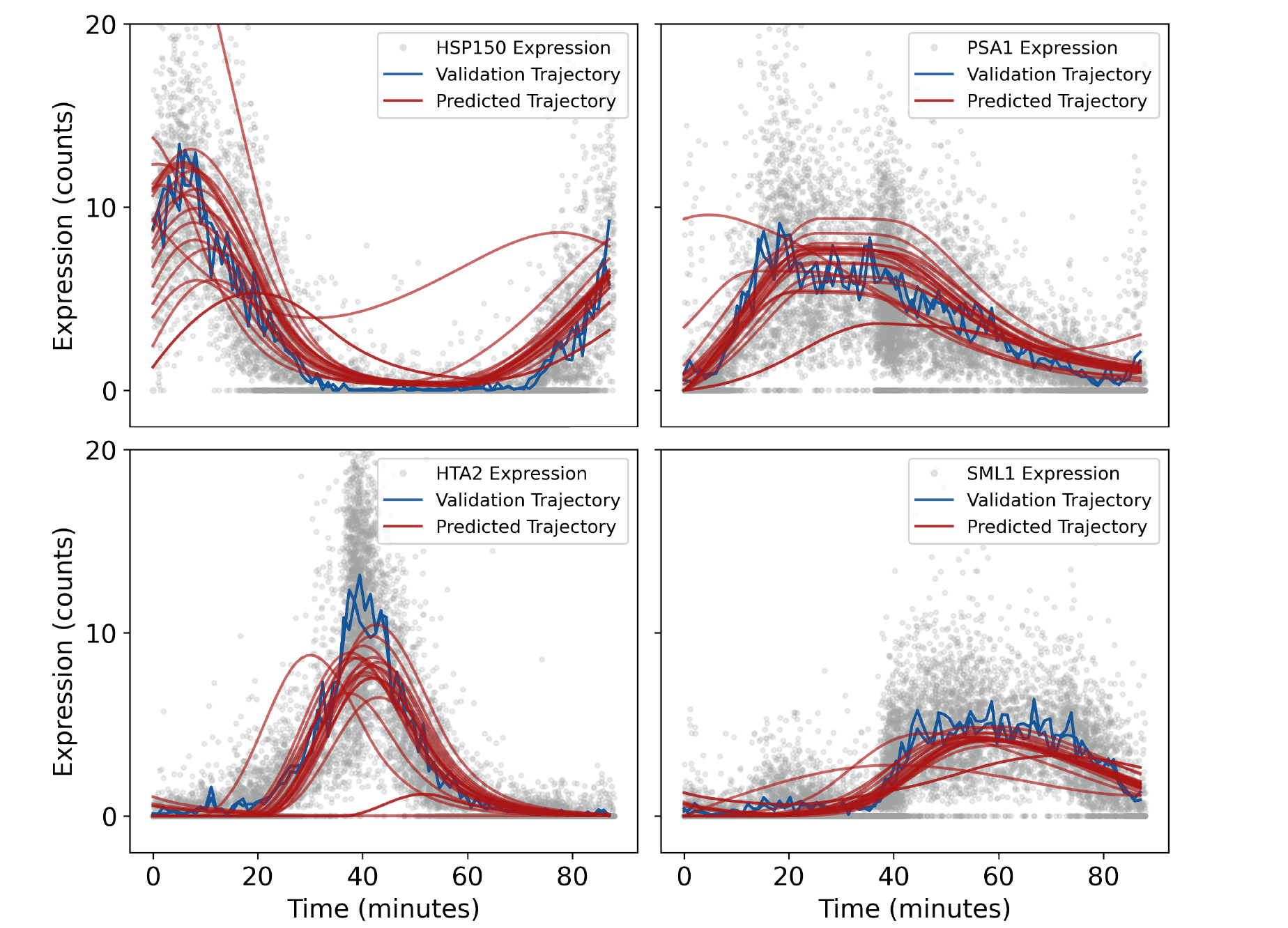
Trajectory Predictions from All Initial Conditions Cell Cycle. Predicted trajectories starting the model from initial conditions outside of the mean trajectory space on validation data. Gray dots are individual validation cells (*n* = 9072); blue lines are the average of 20 cells from 1 minute windows, assembled into a validation trajectory (t=0 to t=88 minutes). Red lines are predicted trajectories (t=0 to t=88 minutes at 1 minute intervals).

**Supplemental Figure S12:**
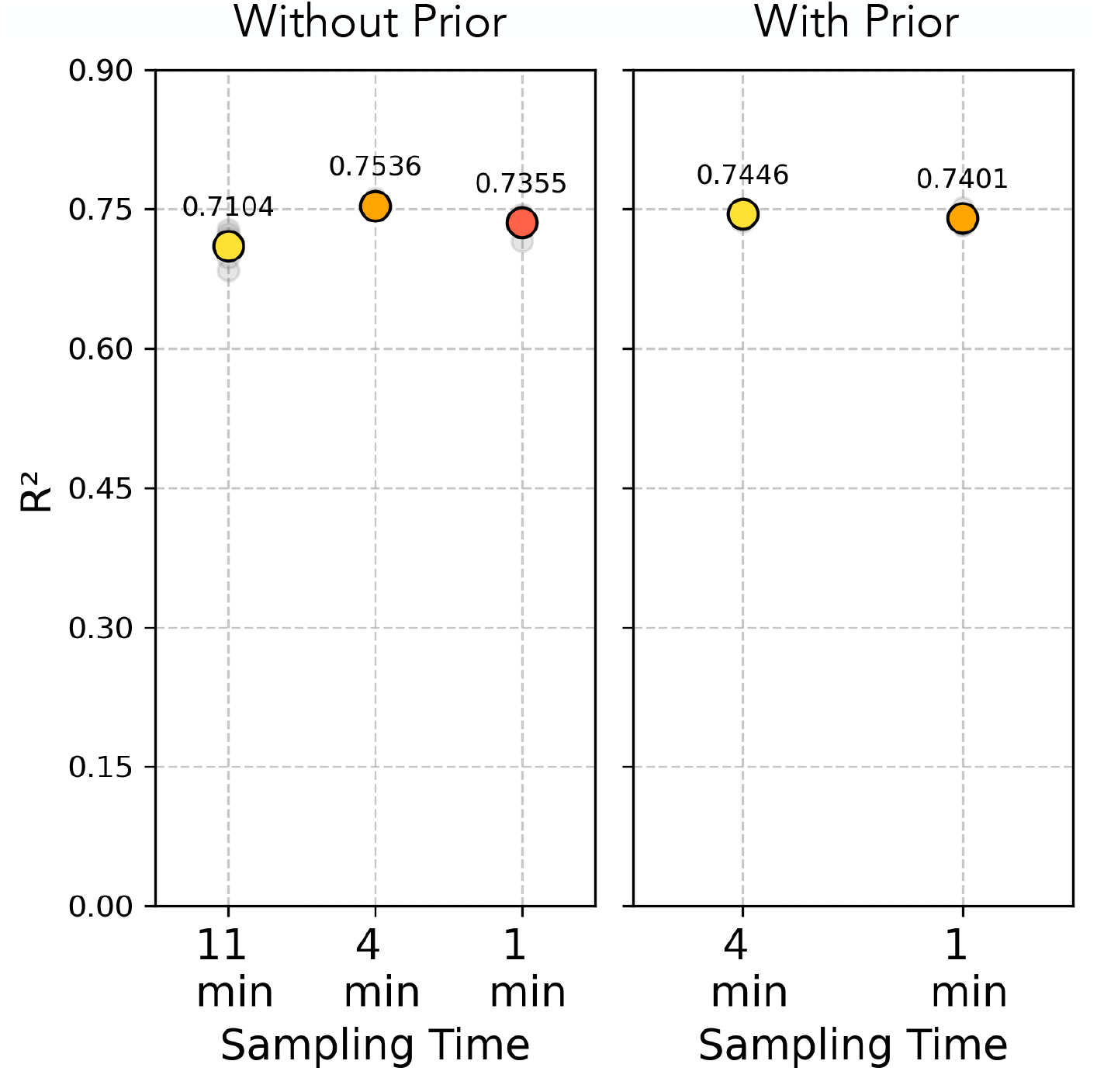
Predictive Capabilities from Sparsely Sampled Trajectories for Cell Cycle. Cells are arranged into trajectories of 88 minutes, with cells sampled every 11 minutes, 4 minutes and 1 minute apart from the trajectory. A scaling stochastic term (see Methods) based on sampling distance was added around each sample time-bin to avoid overfitting. The neural ODE failed to converge on trajectories sampled sparser than 11 minutes without the prior, and sparser than 4 minutes with the prior. However, the R2 values remain high, suggesting the model can still make predictions on datasets less continuously sampled than ours, and that the model can smoothly interpolate predictions at unseen times.

**Supplemental Figure S13:**
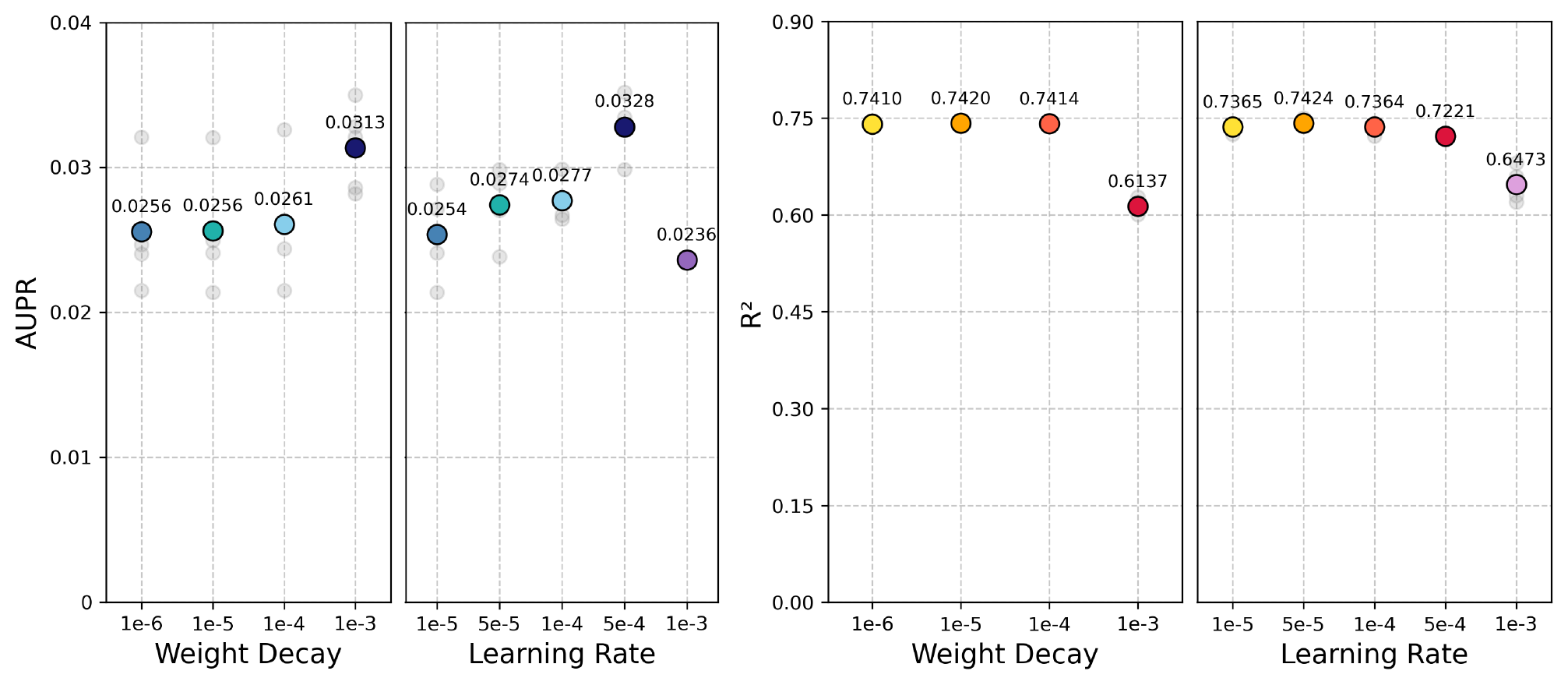
Hyperparameter Grid Search for Cell Cycle. Individual cells are arranged into trajectories of 88 minutes with cells selected one-minute apart. Performance quantified by AUPR and R2 across 5-cross validations. Performance is shown for a range of Adam optimizer learning rates (*γ*) with weight decay (*λ*) held constant at 1*e*^−^5, and a range of weight decays with learning rate held constant at 1*e*^−^4.

**Supplemental Figure S14:**
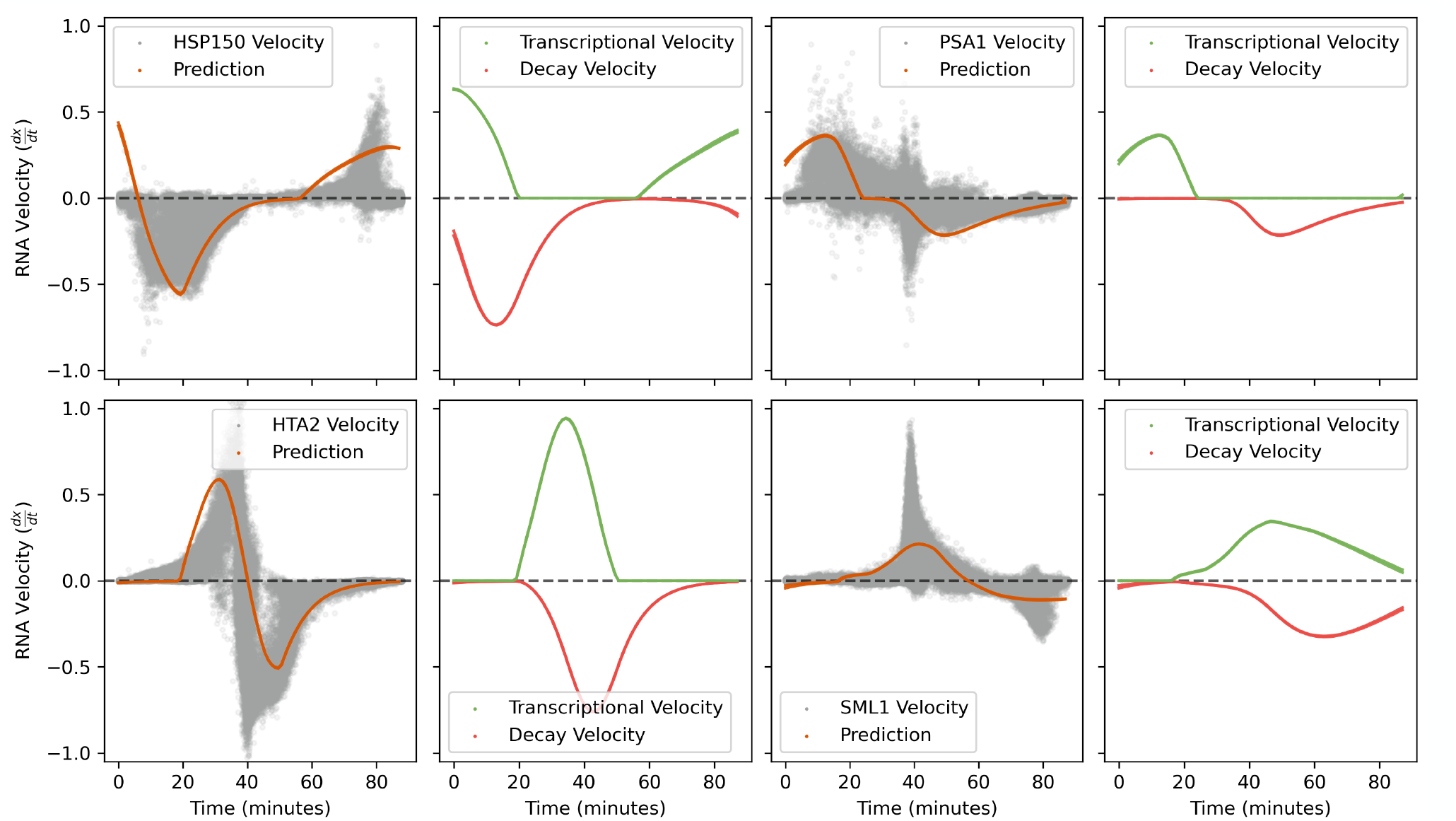
Predicted Biophysical Estimates for Cell Cycle Transcription Factors. Gray dots are estimates of RNA velocity (^*dx*^*/*_*dt*_) for each cell held out as validation, generated by regressing gene expression *x* against time *t* within a local graph neighborhood. Orange lines are in-CAHOOTTS predicted velocities plotted as trajectories. Phase-specific genes *HSP150* (M/G1), *PSA1* (G1), *HTA2* (S), and *CIS3* (G2) are separately plotted. in-CAHOOTTS predicted velocities, separated into green mRNA transcription trajectories (*NN*_*α*_ or 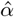) and red mRNA degradation trajectories (− \**x NN*_*λ*_ or 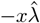.

**Supplemental Figure S15:**
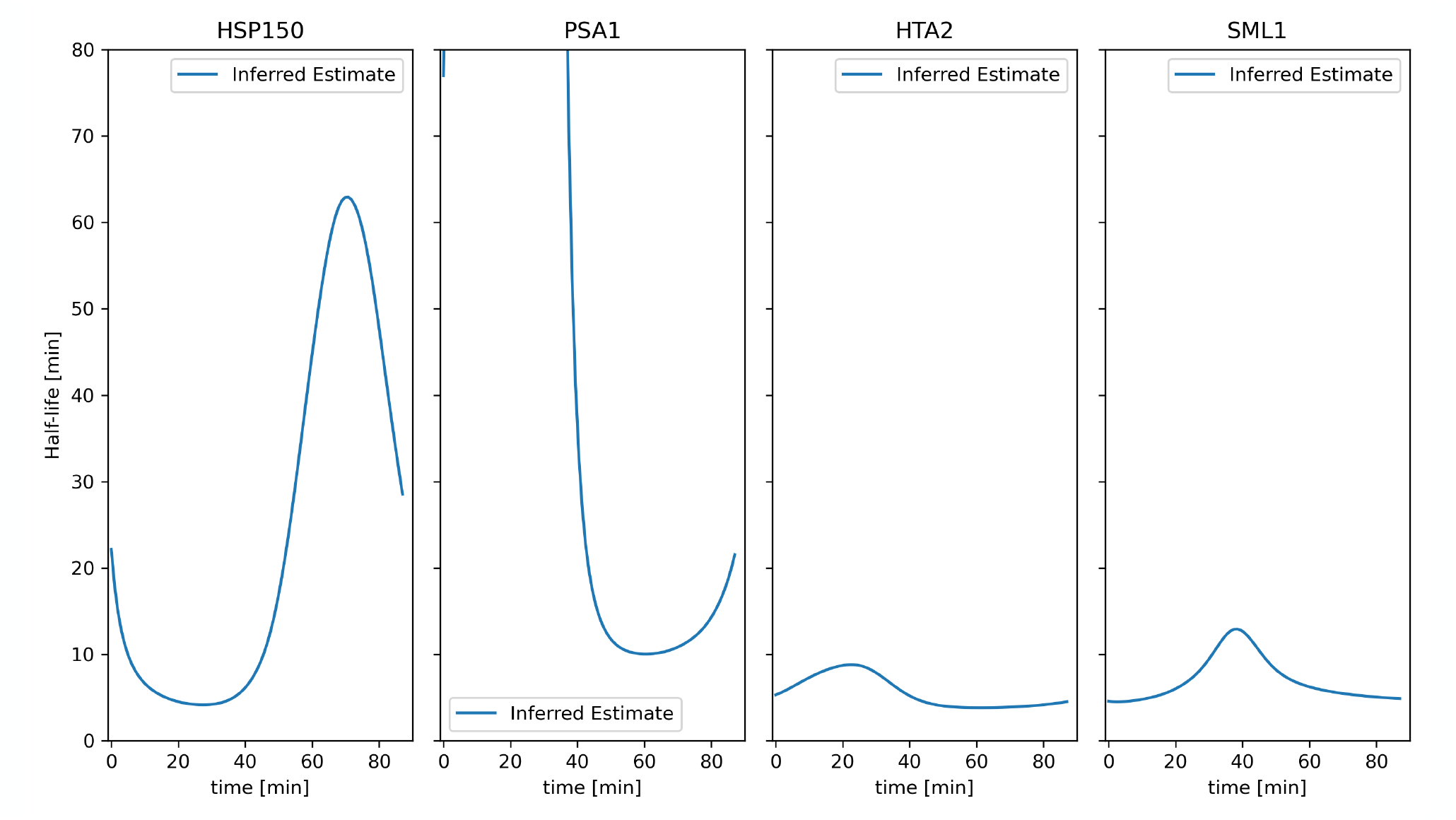
Predicted Half-Lives for Cell Cycle Transcription Factors. in-CAHOOTTS predicted, time-dependent estimate of RNA decay rate trajectories 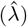 plotted as half-life (*t*_1*/*2_ = ^*log*(2)^*/*_*λ*_). Phase-specific genes *HSP150* (M/G1), *PSA1* (G1), *HTA2* (S), and *CIS3* (G2) are separately plotted.

**Supplemental Figure S16:**
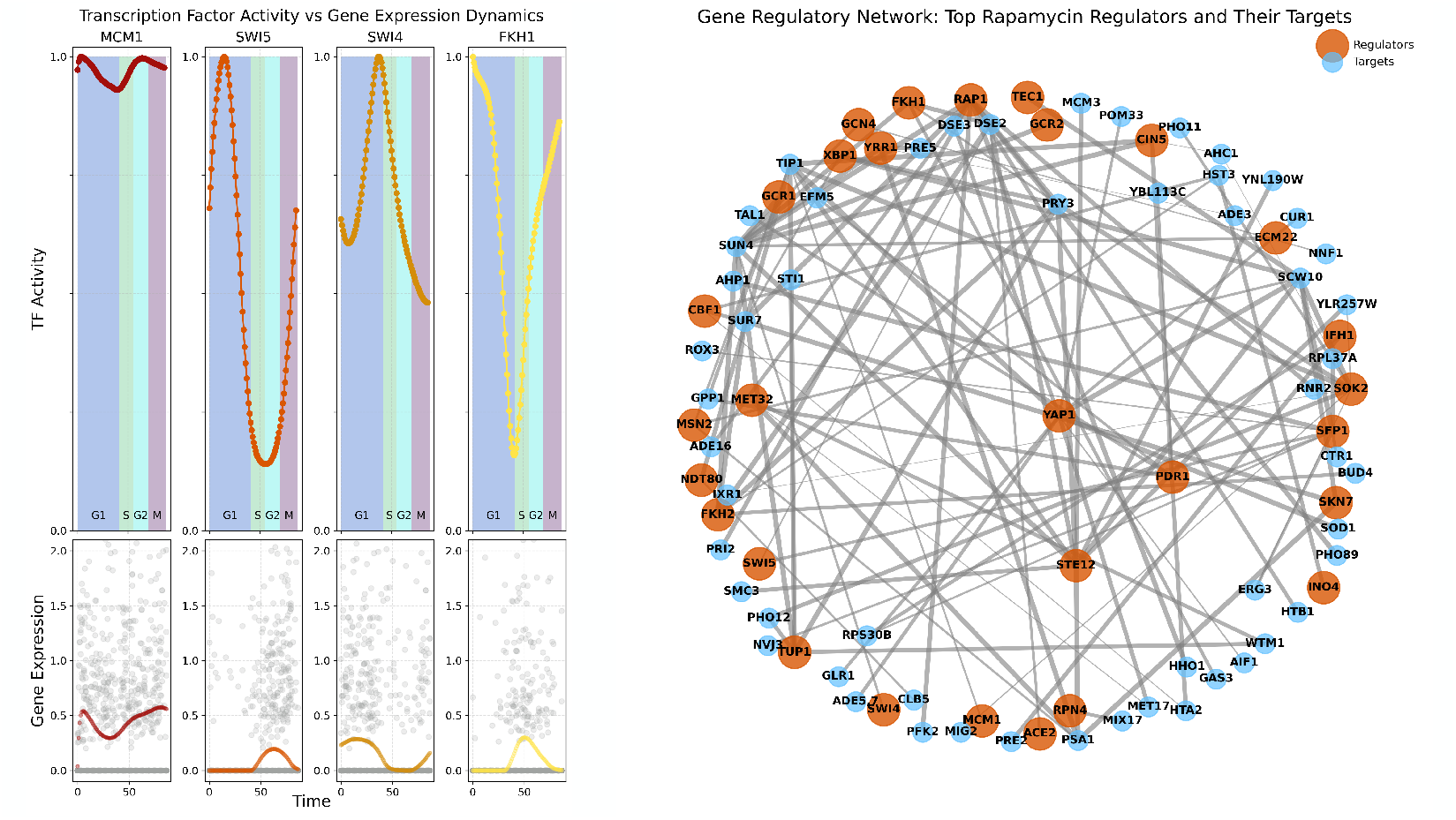
in-CAHOOTTs predicts oscillating TFA and uncovers GRN for cell-cycle dynamics. Predicted transcription factor activities (TFA, top panels) for MCM1, SWI5, SWI4, and FKH1 show distinct temporal patterns across the cell cycle phases [G1 (blue), S (green), G2 (cyan), M (purple)], with SWI5 peaking during late M/early G1, SWI4 during late G1/early S, and FKH1 during late S/G2/M phases. The model-predicted TFA profiles closely correlate with the corresponding target gene expression patterns (bottom panels), demonstrating that computational inference of transcription factor activities can accurately recapitulate the known regulatory dynamics underlying cell cycle progression. Gene regulatory network inferred by in-CAHOOTTS showing predicted cell-cycle transcription factors (orange nodes) and their target genes (blue nodes). Network connections represent high-confidence regulatory relationships learned by the model through explained relative variance analysis, constrained to only connections actually existing within the gold standard gene regulatory network. Gray edges indicate regulatory interactions constrained by prior biological knowledge, demonstrating how the model recovers known regulatory architecture while discovering context-specific activation patterns during rapamycin treatment.

